# Myelin maintenance and addition regulate synaptic plasticity in the adult mouse cortex

**DOI:** 10.64898/2025.12.18.695322

**Authors:** Renee E Pepper, Kalina Makowiecki, Kimberley A Pitman, Raphael Ricci, Angela Nicoli, Phuong Tram Nguyen, Catherine A Blizzard, Ben Emery, Carlie L Cullen, Kaylene M. Young

## Abstract

Myelination of the developing central nervous system (CNS) increases action potential conduction velocity but can also act as a plasticity brake, limiting neurite reorganisation and synaptic plasticity. As myelination is a life-long process, the myelin laid down in development or adulthood could also affect synapse number or plasticity across the lifespan. We report that conditionally deleting the transcription factor, *myelin regulatory factor* (*Myrf*), to disrupt the myelin maintenance program in oligodendrocytes (OLs) from P57 (*Plp-CreER^T2^ :: Myrf ^fl/fl^* mice), led to significant myelin loss and impaired action potential conduction and gross motor performance. By sparsely labelling layer V pyramidal neurons in the primary motor cortex, we could visualise the apical and basal dendrites and their excitatory post-synaptic dendritic spines and determined that spine density was normal in P57+60 *Plp-CreER^T2^ :: Myrf ^fl/fl^* mice. However, a higher proportion of the dendritic spines were large, stable mushroom spines and generated larger amplitude miniature excitatory post-synaptic currents. These data indicate that the myelin maintenance program is critical for preserving neuron adaptability, homeostasis and glutamate sensitivity. When we instead prevented the addition of new OLs and myelin from P57, action potential conduction velocity and gross motor performance appeared normal. Spine density was also normal in *Pdgfrα-CreER^TM^ :: Myrf ^fl/fl^* mice, however, spine morphology was changed along the basal dendrites of layer V M1 pyramidal neurons. At P117, the basal dendrites were frozen in a state that resembled P57 dendrites, as they retained a higher proportion of thin, plastic spines. These data suggest that adult myelination supports the life-long accumulation of stable spines within the basal but not apical dendritic compartment and have important implications for understanding dendritic plasticity rules in the motor circuit.

**Graphical Abstract:** 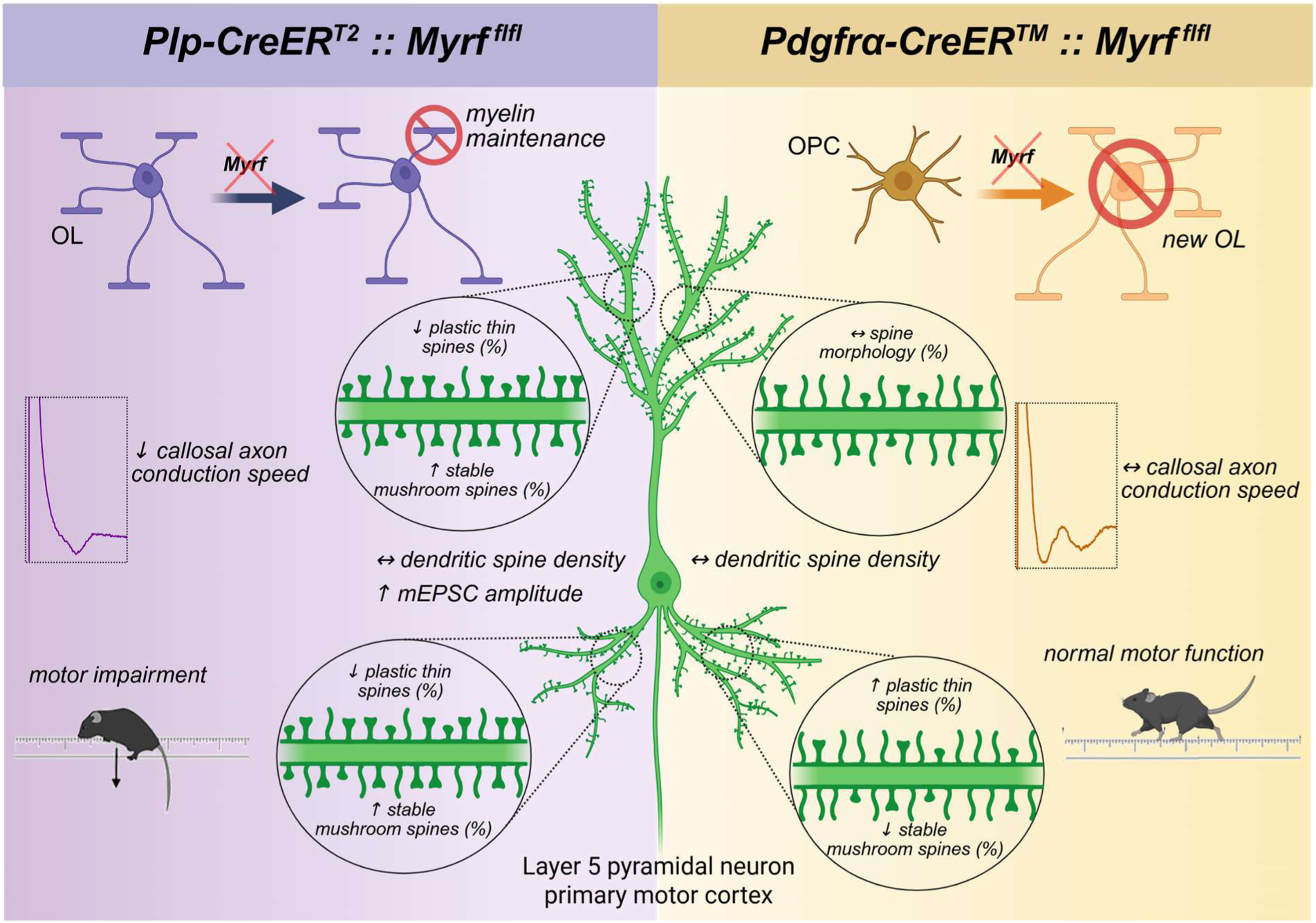

**Highlights:**

- Adult myelin maintenance regulates synaptic plasticity and neuron homeostasis
- Existing myelin regulates layer V pyramidal neuron glutamate sensitivity
- Oligodendrocytes born in young adulthood undergo age-related loss
- New myelin stabilises synapses in the basal but not apical dendritic compartment

**eTOC blurb:** Myelin regulates axon branching, metabolism and action potential conduction speed. We report that myelin also preserves neuron adaptability, homeostasis and glutamate sensitivity in the adult motor cortex. Furthermore, new myelin allows the accumulation of stable synapse on basal but not apical layer V pyramidal neuron dendrites over time.

## Introduction

Oligodendrocytes (OLs) myelinate excitatory ^1–3^ and inhibitory ^4–6^ neurons within the adult central nervous system (CNS), including neurons critical for motor leaning and the precise execution of learned and voluntary movement ^7–10^. Myelination supports the rapid and saltatory conduction of action potentials ^11–13^ and promotes axon health through the provision of metabolites for local ATP synthesis ^14–16^ and the release of exosomes to promote active transport ^17^. In development, cortical myelination coincides with the closure of critical periods when the cortical circuitry and cortical synaptic connections can change rapidly in response to environmental input ^18,19^. Cortical myelination can act as a physical and molecular plasticity brake, as myelination can limit neurite reorganisation ^20^, modulate neurotransmitter release ^21–23^, reduce synaptogenesis ^24^ and synapse turnover ^25^, and regulate the intrinsic excitability of axons ^26^. As OLs are long-lived cells ^3,27^, their myelin has the potential to regulate synaptic plasticity throughout life.

The majority of OLs present in the mature CNS correspond to those generated during development ^27,28^, however, new OLs and myelin are generated throughout life ^2,3,28–32^ to support the long-term recollection of acquired skills and memories ^7,10,33,34^. As memory formation and recall is encoded through synaptic plasticity ^35–37^, this suggests that adult myelination has the capacity to alter synapse number or structure. The acquisition and execution of learned and voluntary movement relies on synaptic plasticity in the primary motor cortex (MC), including those that convey sensory, state and motor information to be the large layer V glutamatergic output neurons ^36–45^. While increasing the activity of layer V pyramidal neurons can increase new OL and myelin addition to the deep layers of the ipsilateral motor cortex and corpus callosum ^8^, it is unclear whether adult myelination modulates the number or stability of intracortical or thalamocortical excitatory inputs to the layer V MC output neurons.

Herein, we show that genetically preventing myelin maintenance by OLs slows compound action potential conduction, impairs motor performance and significantly increases the proportion of spines that are stable mushroom spines along the apical and basal dendrites of layer V MC neurons. We find that myelin maintenance is essential for neuronal homeostasis, and the ability of neurons to balance synaptic plasticity with synaptic stability. By contrast, preventing the addition of new OLs did not alter compound action potential conduction or the execution of previously acquired motor skills. However, it significantly impaired the stabilisation of excitatory synapses on layer V basal but not apical dendrites. Without adult myelination, the basal dendrites of P117 mice appeared “frozen” in a P57 state, with an increased proportion of transient thin spines and fewer stable mushroom spines, suggesting that adult myelination supports the selective and cumulative stabilisation of synapses across the lifespan.

## Results

### In adulthood, conditionally deleting Myrf from OLs prevents myelin maintenance and slows compound action potential conduction

Within mature OLs, the transcription factor, MyRF, supports myelin maintenance ^46,47^. To learn how the myelin maintenance program influences neuronal plasticity, we conditionally deleted *Myrf* from OLs in young adult mice. Tamoxifen was delivered to P57 control (*Myrf ^fl/fl^*) and *Plp-CreERT :: Myrf ^fl/fl^* (referred to as PLP*Myrf ^fl/fl^*) transgenic mice, and brain tissue collected 56 days later (P57+56). Immunohistochemistry, performed on coronal brain sections, revealed that platelet-derived growth factor receptor α (PDGFRα)^+^ OPCs and aspartoacylase (ASPA)^+^ mature OLs were present in the MC (**Fig. 1A, B**) and corpus callosum (CC; **Fig. 1C, D**) of control and PLP*Myrf ^fl/fl^* mice. OPC density was slightly elevated in the PLP*Myrf ^fl/fl^* CC (**Fig. 1E**), consistent with an OPC response to white matter injury ^48^. While ASPA^+^ OL density was normal in the MC and CC of P57+56 PLP*Myrf ^fl/fl^*mice (**Fig. 1F**), staining additional tissue sections to detect: myelin basic protein (MBP) as a proxy for gross myelin content; microglia (ionized calcium-binding adapter protein 1; Iba1), and astrocytes (glial fibrillary acidic protein; GFAP) (**Fig. 1G, H**), revealed that MBP-labelling was reduced (**Fig. 1I**), and a greater area of the CC was covered by Iba1^+^ (**Fig. 1J**) and GFAP^+^ (**Fig. 1K**) pixels. The marked loss of myelin and associated gliosis indicate that conditionally deleting *Myrf* compromised myelin maintenance within 8 weeks but did not cause overt OL loss, consistent with a previous report ^47^.

**Figure 1:**
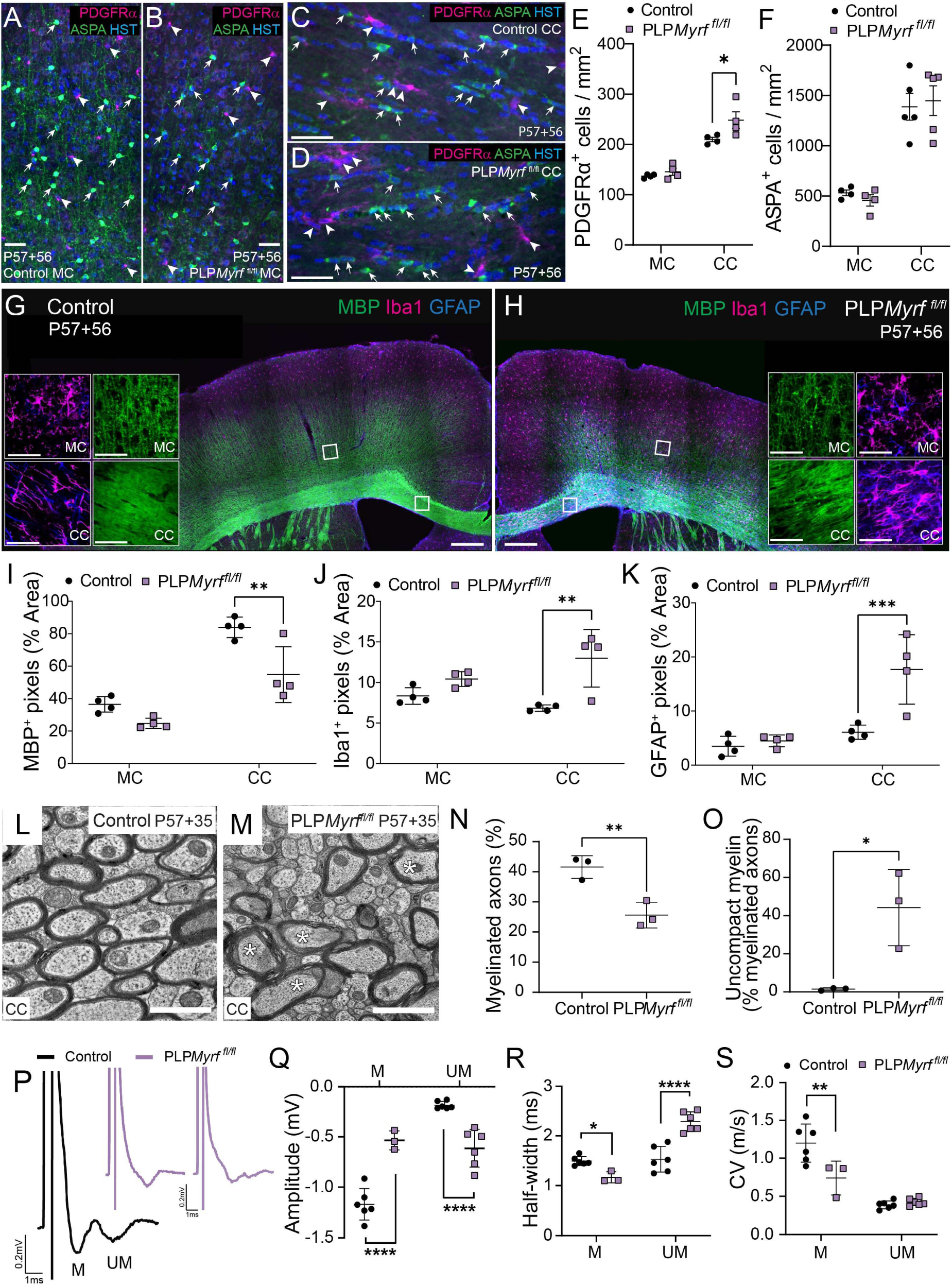
Conditionally deleting Myrf from OLs at P60 results in myelin loss and impaired action potential conduction. (A-D) PDGFRα (magenta) and ASPA (green) labelling in the motor cortex (MC) and corpus callosum (CC) of control (A, C) and PLP*Myrf ^fl/fl^* (B, D) mice at P57+56. Scale bar represents 30µm (A-B) and 40µm (C-D). (E) Quantification of PDGFRα+ OPC density in the MC and CC of control (black circles; n=4) and PLP*Myrf ^fl/fl^* (purple squares; n=4) mice at P57+56. [Two-way ANOVA: Interaction F(1, 12)=2.59, p=0.133; region F(1,12)= 89.43, p=<0.0001; genotype F(1,12)=6.44, p=0.026]. (F) Quantification of ASPA+ OL density in the MC (n=4 per group) and CC (n=5 per group) of control and PLP*Myrf ^fl/fl^* mice at P57+56. [Two-way ANOVA: Interaction F(1, 14)=0.34, p=0.568; region F(1,14)= 63.25, p=<0.0001; genotype F(1,14)=0.004, p=0.951]. (G-H) MBP (green), Iba1 (magenta) and GFAP labelling (blue) in the MC and CC of control (G) and PLP*Myrf ^fl/fl^* (H) mice at P57+56. Scale bar represents 200µm and 40µm (inset). (I) Quantification of the proportion of area containing MBP+ pixels in the MC and CC of control (n=4) and PLP*Myrf ^fl/fl^* (n=4) mice at P57+56. [Two-way ANOVA: Interaction F(1, 12)=3.26, p=0.096; region F(1,12)= 65.16, p=<0.0001; genotype F(1,12)=18.04 p=0.0011]. (J) Quantification of the proportion of area containing Iba1+ pixels in the MC and CC of control (n=4) and PLP*Myrf ^fl/fl^* (n=4) mice at P57+56. [Two-way ANOVA: Interaction F(1,12)=4.49, p=0.055; region F(1,12)=0.31, p=0.58; genotype F(1,12)=18.48 p=0.0010]. (K) Quantification of the proportion of area containing GFAP+ pixels in the MC and CC of control (n=4) and PLP*Myrf ^fl/fl^* (n=4) mice at P57+56. [Two-way ANOVA: Interaction F(1,12)=9.52, p=0.009; region F(1,12)=21.08, p=0.0006; genotype F(1,12)=13.43 p=0.0032]. (L-M) Transmission electron micrographs from the CC of P57+35 control and PLP*Myrf ^fl/fl^* mice. Scale bar represents 1µm. (N) Quantification of the proportion of myelinated axons in the CC of control (n=3) and PLP*Myrf ^fl/fl^* (n=3) mice at P57+35. [Unpaired t-test: t=4.85, df=4; p=0.0083] (O) Quantification of the proportion of myelinated callosal axons with loose, uncompact myelin in control (n=3) and PLP*Myrf ^fl/fl^* (n=3) mice at P57+35. [Unpaired t-test: t=3.703, df=4; p=0.0208] (P) Example CAP traces from the CC of P57+35 control (black) and PLP*Myrf ^fl/fl^* (purple) mice. (Q) Amplitude (mV) of the myelinated (M) and unmyelinated (UM) peaks of callosal CAPs measured in P57+35 control (n=6) and PLP*Myrf ^fl/fl^* (n=6) mice. [Two-way ANOVA: Interaction F(1,17)=71.70, p<0.0001; peak F(1,17)=52.19, p<0.0001; genotype F(1,17)=2.61 p=0.124]. (R) Half-width (ms) of the myelinated (M) and unmyelinated (UM) peaks of callosal CAPs measured in P57+35 control (n=6) and PLP*Myrf^fl/fl^* (n=6) mice. [Two-way ANOVA: Interaction F(1,17)=40.36, p<0.0001; peak F(1,17)=55.89, p<0.0001; genotype F(1,17)=6.42 p=0.021]. (S) Conduction velocity for myelinated (M) and unmyelinated (UM) axons measured from callosal CAP traces recorded from P57+35 control (n=6) and PLP*Myrf^fl/fl^* (n=6) mice. [Two-way ANOVA: Interaction F(1,17)=11.48, p=0.0035; peak F(1,17)=58.48, p<0.0001; genotype F(1,17)=8.28 p=0.01]. Graphs show mean ± SD. *p<0.05; **p<0.01; ***p<0.001; ****p<0.0001 by Bonferroni’s post-test or t-test (N-O).

To more precisely evaluate the extent to which myelin maintenance was compromised, we performed transmission electron microscopy (TEM) at an earlier timepoint. Even at P57+35, fewer CC axons were myelinated in PLP*Myrf ^fl/fl^* mice (**Fig. 1L-N**) and a higher proportion of the myelinated axons showed evidence of myelin decompaction (**Fig. 1O**). When we measured compound action potential (CAPs) conduction across the CC in acute brain slices from P57+35 control mice, we consistently observed the two peaks (**Fig. 1P**) that correspond to action potentials conducted along myelinated and unmyelinated axons ^49^. By contrast, CAPs recorded from PLP*Myrf ^fl/fl^* brain slices had a smaller or absent myelinated axon peak (**Fig. 1P**). When present, the amplitude and half-width of the myelinated axon peak was significantly reduced, while the amplitude and half-width of the unmyelinated axon peak was increased (**Fig. 1Q**, **R**), indicating that a greater proportion of axons were unmyelinated in the PLP*Myrf ^fl/fl^* CC. As myelinated CAP velocity was also reduced (**Fig. 1S**), these data indicate that disrupting the myelin maintenance program results in functional demyelination by P57+35.

### The adult deletion of Myrf from OLs leads to motor dysfunction

When myelin maintenance is compromised in the CNS, behavioral modalities, including motor coordination and performance, can be impaired ^47^. Therefore motor performance was evaluated at P56, one day prior to tamoxifen delivery, and then weekly for 6-8 weeks (up to P57+56). Mice were placed on the rotarod to detect large changes in motor circuit and / or muscle function, such as those seen in clinical models of amyotrophic lateral sclerosis ^50^. The rotarod performance of PLP*Myrf ^fl/fl^* mice appeared poor relative to controls, but this did not reach statistical significance at any individual timepoint (**Fig. 2A**). By contrast, when mice were placed on the grid-walk to evaluate dextrous fine motor skill, we found that the performance of PLP*Myrf ^fl/fl^* mice was significantly impaired by P57+35 (**Fig. 2B**). Similarly, control mice consistently made few beam-walk foot-slip errors across the time course, while the performance of PLP*Myrf ^fl/fl^* mice had declined significantly by P57+21 (**Fig. 2C**).

**Figure 2:**
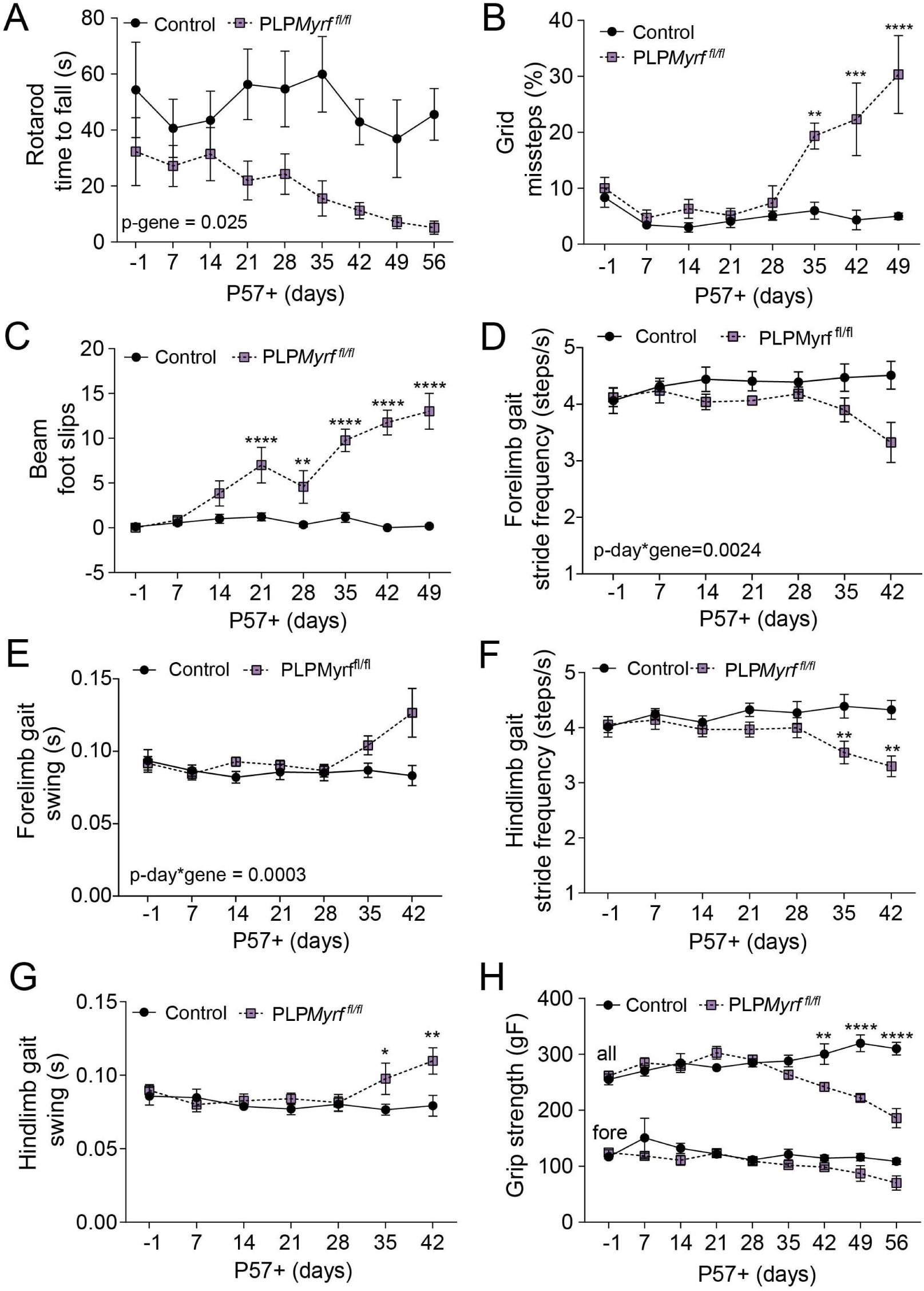
The conditional deletion of Myrf from OLs at P60 impairs motor function within 21 days. (A) Time taken for control (black circles) and PLP*Myrf ^fl/fl^* (purple squares) mice to fall from the rotarod at P57-1 to P57+56. [Restricted Maximum Likelihood (REML) mixed effects model: Interaction F(8, 84)=0.801 p=0.603; day F(2.86, 30.04)=1.73 p=0.18; genotype F(1,14)=6.31 p=0.025]. (B) Proportion of missteps made by control and PLP*Myrf ^fl/fl^* mice performing the grid walk test at P57-1 to P57+49. [Restricted Maximum Likelihood (REML) mixed effects model: Interaction F(7, 64)=7.601 p<0.0001; day F(2.52, 23.05)=8.626 p=0.0008; genotype F(1,14)=27.26 p<0.0001]. (C) Number of foot slips made by control and PLP*Myrf ^fl/fl^* mice performing the beam walk test at P57-1 to P57+49. [Restricted Maximum Likelihood (REML) mixed effects model: Interaction F(7,74)=14.2 p<0.0001; day F(1.7, 18.2)=14.65 p=0.0003; genotype F(1,14)=91.73 p<0.0001]. (D-E) Average forelimb stride frequency (D) and limb swing time (E) of control and PLP*Myrf ^fl/fl^* mice during treadmill (DigiGait^TM^) running from P57-1 to P57+42. [Restricted Maximum Likelihood (REML) mixed effects model: *Stride frequency* (Interaction F(6, 52)=4.41 p=0.0011; day F(2.99, 25.94)=1.53 p=0.229; genotype F(1,12)=3.81 p=0.074); *Swing* (Interaction F(6, 53)=5.24 p=0.0003; day F(1.930, 17.05)=2.47 p=0.116; genotype F(1,13)=3.145 p=0.099)] (F-G) Average hindlimb stride frequency (F) and limb swing time (G) of control and PLP*Myrf ^fl/fl^* mice during treadmill (DigiGait^TM^) running from P57-1 to P57+42. [Restricted Maximum Likelihood (REML) mixed effects model: *Stride frequency* (Interaction F(6, 52)=4.41 p=0.0011; day F(2.99, 25.94)=1.53 p=0.229; genotype F(1,12)=3.81 p=0.074); *Swing* (Interaction F(6, 52)=5.42 p=0.0002; day F(2.02, 17.48)=3.06 p=0.072; genotype F(1,12)=1.68 p=0.21)] (H) Grip strength for all paws and forepaws only of control and PLP*Myrf ^fl/fl^* mice from P57-1 to P57+56. [Restricted Maximum Likelihood (REML) mixed effects model: *All paws* (Interaction F(8, 84)=14.27 p<0.0001; day F(3.16, 33.25)=4.37 p=0.0096; genotype F(1,14)=10.02 p=0.0069); *Fore paws* (Interaction F(8, 84)=0.79 p=0.62; day F(1.57, 16.50)=2.08 p=0.16; genotype F(1,14)=4.18 p=0.06)] Graphs show mean ± SEM. Control n= 6-9; PLPMyrf *^fl/fl^* n= 4-7. *p<0.05; **p<0.01; ***p<0.001; ****p<0.0001 by Bonferroni’s post-test or t-test. See also Figure S1

We predicted that preventing OL myelin maintenance would also disrupt established motor patterns, like those that underpin gait, and found that gait was impacted by P57+35 (**Fig. 2D-G** and **Fig. S1**). While forelimb gait parameters were largely unaffected over the testing period (**Fig. 2D-E**; **Fig. S1**), PLP*Myrf ^fl/fl^* hindlimb stride frequency was reduced by P57+35 **(Fig. 2F**) and gait swing time was increased (**Fig. 2G**). These data are consistent with previous reports of ataxia following *Myrf* deletion ^47,51^ and suggest that the relative movement of fore and hindlimbs becomes uncoordinated. The majority of PLP*Myrf ^fl/fl^*mice (∼70%) failed to walk on the Digigait treadmill after P57+42, preventing further reliable recordings from being obtained. As PLP*Myrf ^fl/fl^* forepaw grip strength was normal, but the grip strength of all four paws was significantly weaker from P57+42 (**Fig. 2H**), myelin may be more rapidly compromised in circuits that regulate hind paw movement or the ability of mice to coordinate movement of the fore and hind paws.

### Myelin maintenance allows layer V pyramidal neurons to retain a high proportion of thin, highly plastic, dendritic spines

The MC is particularly critical for the learning and execution of dexterous motor skills ^36,43,44^. The major output neurons, the layer V pyramidal neurons extend their apical dendrite dorsally to form tuft dendrites and receive input from layer 1, and directly elaborates basal dendrites from the soma to receive input within layer V ^52,53^. The apical and basal dendritic compartments are functionally distinct ^54^ and allow MC excitatory neurons to receive and integrate cortico-cortical and thalamo-cortical inputs ^38^. The spines protruding from excitatory neuron dendrites correspond to the postsynaptic compartment of glutamatergic synapses and transition from small thin spines to larger mushroom spines as they are stabilised and strengthened through long-term potentiation ^55,56^. In the developing visual cortex, myelination decreases the density of spines along layer V pyramidal neuron dendrites ^25^.

### However, it is unclear whether the OL myelin maintenance program impact synaptic plasticity in adulthood

To determine whether the myelin laid down before 2 months of age, has an ongoing neuronal regulatory function that impacts dendritic spine density or structural plasticity, we crossed the PLP*Myrf ^fl/fl^* with *Thy1-YFPH* transgenic mice. The *Thy1-YFPH* transgene drives YFP expression in a subset of MC pyramidal neurons, supporting visualisation of the apical and basal dendrites and dendritic spines (**Fig. 3A-B**). Individual YFP^+^ pyramidal neurons were identified in layer V of the MC of P57+60 *Thy1-YFPH* control and PLP*Myrf ^fl/fl^* mice, and high-resolution confocal images collected along the apical and basal dendrites for 3D reconstruction (**Fig. 3C, D**). The overall density of dendritic spines was ∼19 spines / 10µm length of apical or basal dendrite and was equivalent in control and PLP*Myrf ^fl/fl^* mice (**Fig. 3E**). As spines undergo structural changes that reflect their functional state, we next classified each spine into one of three morphological types: thin, stubby or mushroom (**Fig. 3B-D**). The apical dendrites of PLP*Myrf ^fl/fl^*mice had an increased density of stable mushroom spines compared to control mice (**Fig. 3F**). The density of mushroom spines was similarly increased along the basal dendrites of PLP*Myrf ^fl/fl^* mice, at the expense of thin, plastic spines (**Fig. 3F**).

**Figure 3:**
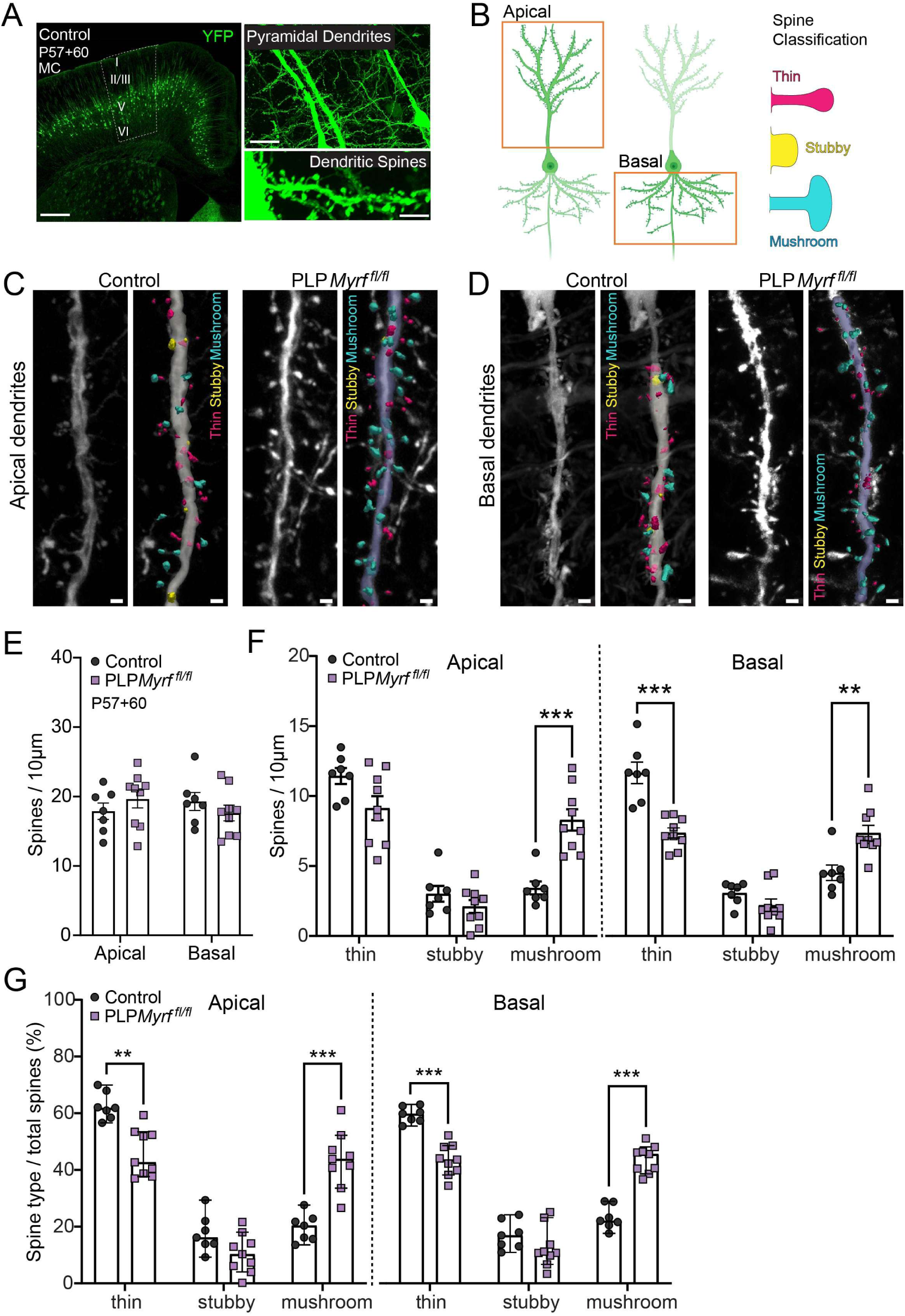
At P57, the conditional deletion of Myrf from oligodendrocytes promotes the morphological plasticity of layer V pyramidal neuron dendritic spines, and the generation of stable mushroom spines. (A) YFP (green) labelling of layer V pyramidal neurons, including dendrites and dendritic spines in the motor cortex of P57+60 *Thy1-YFPH* mice. Scale bars represent 200µm (left), 40µm (top right), 5µm (bottom right). (B) Schematic representation of apical and basal dendrites of cortical layer V pyramidal neurons and thin (magenta), stubby (yellow) and mushroom (cyan) spine type classification. (C-D) Representative images and rendered traced segments of Thy1YFP+ layer V pyramidal neuron apical (A), and basal (B) dendrites, from control and PLPMyrf^fl/fl^ mice (P57+60). Rendered images show example spines classified by morphology as ‘thin’ (magenta), stubby (yellow) or mushroom (cyan) subtypes. Scale bars represent 1μm. (E) Mean dendritic spine density (number of spines per 10µm dendrite length), in apical and basal dendrites. Overall spine density did not differ significantly between genotypes in apical or basal dendrites [2-way mixed ANOVA; genotype x dendrite interaction: F(1,14) = 2.533, p= 0.134; genotype (control, PLPMyrf^fl/fl^): F (1, 14) = 0.104, p = 0.751; dendrite (within-subjects: apical, basal): F(1,14) = 0.237, p=0.634]. (F) Mean spine densities (number spines per 10µm dendrite length) of each morphological subtype (thin, mushroom, stubby), for apical and basal dendrites. Bars = means, error bars: SEM. Densities of subtypes were significantly different between control and PLPMyrf^fl/fl^ mice [3-way mixed ANOVA run with within-subject factors: dendrite (apical, basal), spine type (thin, mushroom, stubby) and between-subjects factor: genotype (control, PLPMyrf^fl/fl^); 3-way interaction: dendrite*spine type* genotype, F(2,28) = 1.327, p= 0.281); 2-way interactions: spine type* genotype: F(2,28) = 39.56, p<0.001; dendrite*genotype: F(1,14) = 2.516, p = 0.135; dendrite*spine type: F(2,28)=0.915, p = 0.412; dendrite: F(1,14) = 0.249, p = 0.625; spine type: F(2,28) = 159.973, p <0.001; genotype: F(1,14) = 0.09, 0.765]. (G) Dendritic spine morphological classifications as percentage total spines (spine type/ total spines). Bars = median, error bars: 95%CIs. Control vs. PLPMyrf^fl/fl^ mean ranks Mann-Whitney U, Bonferroni-corrected p-values; apical dendrites percent thin: z = -3.123, p = 0.002; apical dendrites percent mushroom: z = 3.228, p = 0.001. Basal dendrites percent thin: z = -3.334, p <0.001; Basal dendrites percent mushroom: z = 3.334, p<0.001. Percentage of spines classified as ‘stubby’ spines did not differ between genotypes in apical or basal dendrites (p-values >0.05). Datapoints in all graphs represent individual mice (mean of ≥ 5 dendrite segments per mouse); asterisks denote Bonferroni-adjusted p-values: *p<0.05, **p<0.01, ***p<0.001.

The maturation of thin spines into mushroom spines is a structural correlate of long-term potentiation, with larger spine heads having a higher concentration of AMPA receptors ^55^. As a predominance of thin spines was replaced by an approximately equal proportion of thin and mushroom spines in PLP*Myrf ^fl/fl^* mice (**Fig. 3G**), we predicted that the layer V MC pyramidal neurons would generate larger miniature-excitatory post synaptic currents (mEPSCs) in response to spontaneous quantal glutamate release. To evaluate this, we generated acute coronal brain slices from P57+65 control or PLP*Myrf ^fl/fl^* mice for whole cell patch clamp electrophysiology. The capacitance and membrane resistance of layer V pyramidal neurons was equivalent in control and PLP*Myrf ^fl/fl^* mice (**Fig. 4A**). When mEPSCs were recorded at the soma (**Fig. 4B**), the average amplitude of mEPSC events was significantly increased in PLP*Myrf ^fl/fl^* mice (**Fig. 4C**). This was associated with a rightward shift in the cumulative distribution plot of mEPSC amplitude (**Fig. 4D**). Released glutamate vesicles producing larger post-synaptic depolarizations is consistent with more spines being mushroom spines (**Fig. 3G**). mEPSC frequency was also elevated in PLP*Myrf ^fl/fl^* mice (**Fig. 4E**), which could reflect an increase in the number of pre-synaptic sites or an increased probability of presynaptic vesicle release in PLP*Myrf ^fl/fl^* mice. Overall, these data indicate that the OL myelin maintenance program influences neuronal homeostasis and glutamate sensitivity in the adult mouse MC.

**Figure 4:**
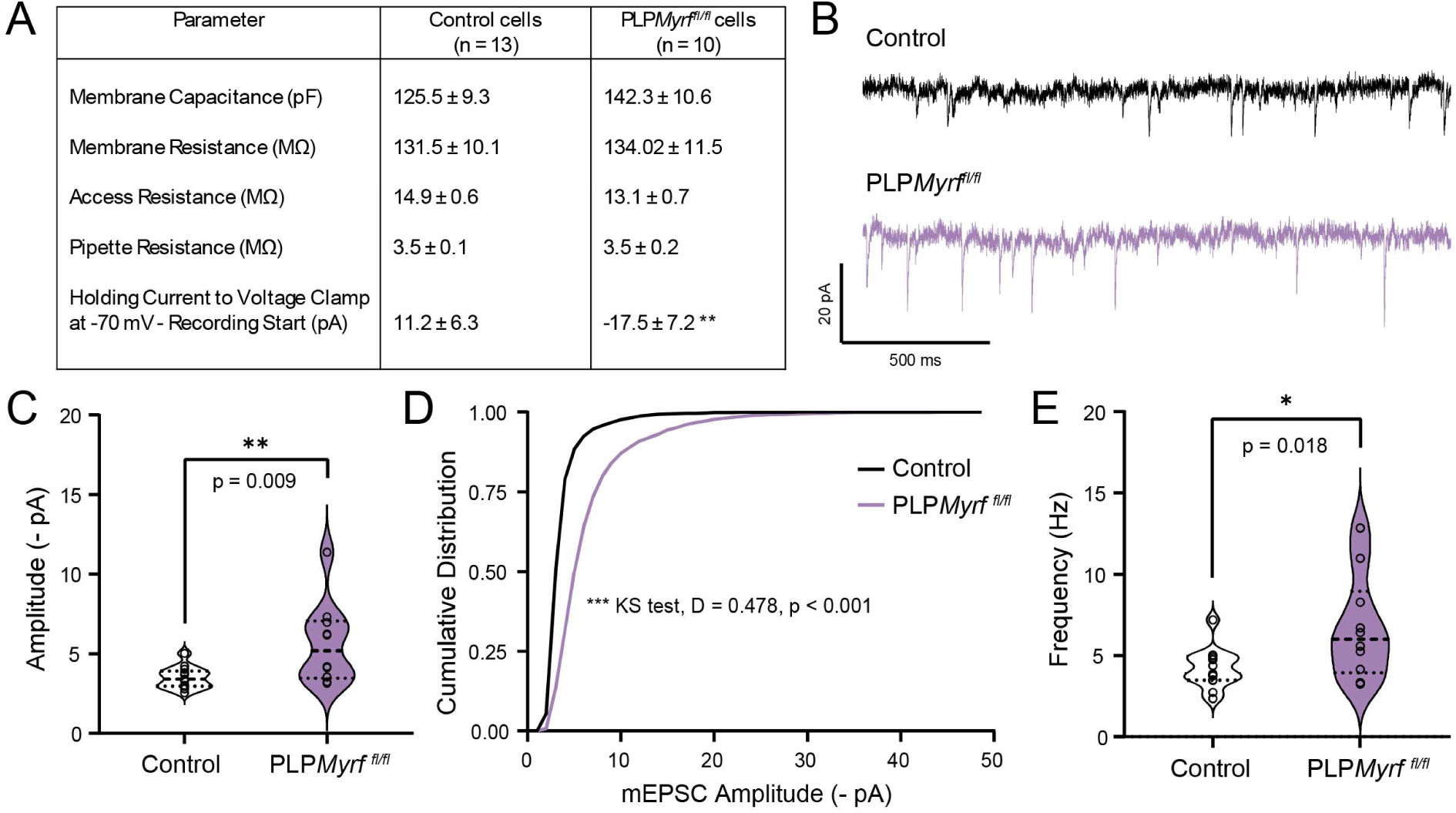
In the motor cortex of P57+65 PLPMyrf ^fl/fl^ mice, layer V pyramidal neurons generate larger mEPSCs in response to vesicular glutamate release. (A) Table of membrane properties and other measurements acquired during the whole cell patch clamp recordings of layer V pyramidal neurons from P57+65 control and Plp*Myrf ^fl/fl^* mice. Parameters were compared between groups using unpaired two-tailed (independent samples) t-tests. (B) Representative trace showing mEPSCs recorded from layer V pyramidal neurons of P57+65 control or Plp*Myrf ^fl/fl^* mice. (C) Average amplitude of mEPSCs recorded from layer V pyramidal neurons of P57+65 control and Plp*Myrf ^fl/fl^* mice. Recordings were taken from n=13 neurons from the slices of n=5 P57+65 control mice, and n=10 neurons from slices from n=3 Plp*Myrf ^fl/fl^* mice. Average amplitude was compared by two-tailed unpaired t-test (t = 2.892), p<0.01. (D) Cumulative distribution plot for mEPSC amplitudes for mEPSCs recorded from layer V pyramidal neurons in P57+65 control and Plp*Myrf ^fl/fl^* mice. The distributions were compared by a two-sample Kolmogorov-Smirnov test: D = 0.478, p<0.001. (E) Average frequency of mEPSCs recorded from the soma of layer V pyramidal neurons of P57+65 control and Plp*Myrf ^fl/fl^* mice. Recordings were taken from n=13 neurons from 5 individual P57+65 control mice, and n=10 neurons from 3 individual Plp*Myrf ^fl/fl^* mice. Average frequency was compared by two-tailed unpaired t-test (t = -2.572), p <0.05. Datapoints in C and E represent individual neurons. Asterisks denote p-values: *p<0.05, **p<0.01, ***p<0.001.

### The conditional deletion of Myrf from adult OPCs prevents further OL addition and myelination

OLs are added to the brain throughout adulthood ^9,30^ and are required for memory consolidation ^33,34^. As memory consolidation requires synaptic plasticity ^57^, we evaluated the capacity of adult-born OLs to influence synapse number or structure, even under basal conditions. To conditionally delete *Myrf* from OPCs in the adult mouse CNS and prevent new OL maturation, tamoxifen was administered to P57 control (*Pdgfrα-CreER^TM^:: Rosa26-YFP)* and Rα*Myrf ^fl/fl^ (Pdgfrα-CreER^TM^ :: Rosa26-YFP :: Myrf ^fl/fl^)* transgenic mice. As new OPCs can be generated from neural stem cells in the adult mouse subventricular zone ^58,59^, tamoxifen was also administered once every 90 days throughout the study, to capture these cells. Immunohistochemistry on coronal brain cryosections detected PDGFRα, OLIG2 and YFP in the MC and CC of control and Rα*Myrf ^fl/fl^* mice from P57+30 to P57+345 (**Fig. 5A-L**). The density of PDGFRα^+^ OPCs was equivalent in control and Rα*Myrf ^fl/fl^* mice, and the recombined YFP^+^ fraction unaffected by genotype (**Fig. S2**). In the MC, control mice had significantly more YFP^+^ MC OLs (PDGFRα-neg OLIG2^+^ YFP^+^) than Rα*Myrf ^fl/fl^* mice by P57+30 (**Fig. 5M**). In the CC of Rα*Myrf ^fl/fl^* mice, the proportion of YFP^+^ cells that were PDGFRα-neg OLIG2^+^ OLs was reduced by ∼59% at P57+30 (**Fig. 5N**). By P57+120, new OL number was reduced by ∼79% in the MC (**Fig. 5M**) and ∼82% in the CC (**Fig. 5N**). This is largely due to the impaired survival of premyelinating OLs, as P57+120 Rα*Myrf ^fl/fl^* mice had ∼76% fewer YFP^+^ BCAS1^+^ (breast carcinoma amplified sequence 1) premyelinating OLs in the MC and ∼84% fewer in the CC than control mice (**Fig. S2**). We also noted that the proportion of YFP^+^ cells that were OLs declined significantly in the CC of control mice between P52+120 and P57+345 (**Fig. 5N**). OL die in older mice ^31^, including within white matter tracts ^60^, and our data may suggest that adult-born CC OLs are highly susceptible to age-related loss.

**Figure 5:**
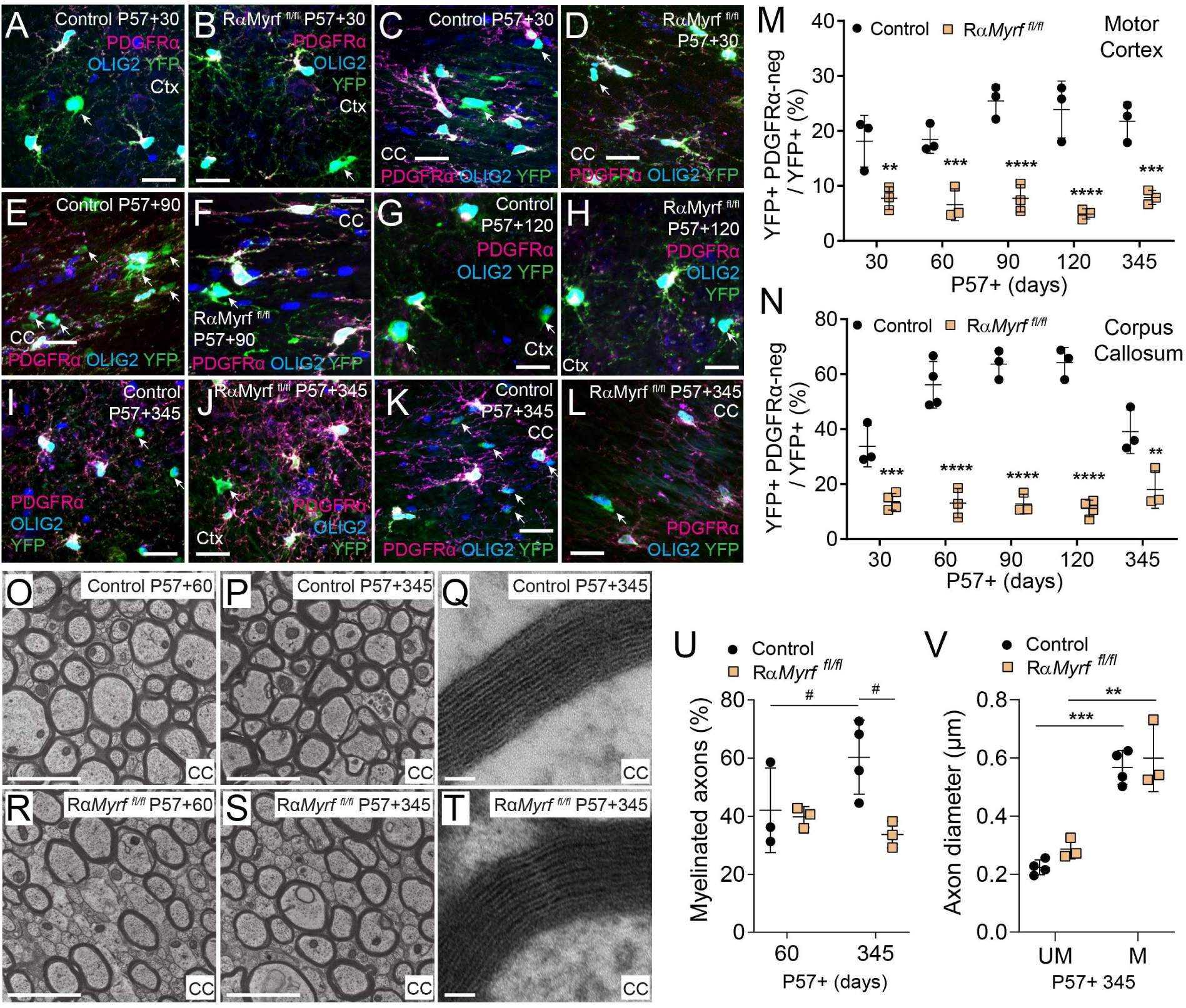
Conditionally deleting Myrf from OPCs at P57, blocks adult oligodendrogenesis and myelination. (A-L) Confocal images from the motor cortex (MC) and corpus callosum (CC) of control and Rα*Myrf ^fl/fl^* mice, stained to detect OPCs (PDGFRα, magenta), cells of the OL lineage (OLIG2, blue) and YFP (green). (M) Quantification of the proportion of YFP+ cells that are PDGFRa-negative OLIG2+ OLs in the MC of control and Rα*Myrf ^fl/fl^* mice between P57+30 and P57+345. [2-way between-subjects ANOVA, timepoint*genotype interaction: F (4, 20) = 2.100, p = 0.1187; timepoint: F (4, 20) = 1.627, p = 0.2066; genotype: F (1, 20) = 163.0, p <0.001]. (N) Quantification of the proportion of YFP+ cells that are PDGFRa-negative OLIG2+ OLs in the CC of control and Rα*Myrf ^fl/fl^* mice between P57+30 and P57+345. [2-way between-subjects ANOVA, timepoint*genotype interaction: F(4,23) = 11.91, p <0.001; timepoint: F (4, 23) = 7.459, p <0.001; genotype: F (1, 23) = 330.0, p <0.001]. (O-T) Transmission electron micrographs through the CC of P57+60 and P57+345 control and Rα*Myrf ^fl/fl^* mice. Scale bars represent 2µm (O-P, R-S), 50nm (Q, T) (U) Proportion of callosal axons that are myelinated in the CC of control and Rα*Myrf ^fl/fl^* mice at P57+60 and P57+345. [2-way ANOVA, timepoint*genotype interaction: F (1, 9) = 4.356, p = 0.0665; timepoint: F (1, 9) = 1.091, p = 0.3235; genotype: F (1, 9) = 6.158, p =0.0349]. (V) The average diameter of unmyelinated (UM) and myelinated (M) axons in the CC of control and Rα*Myrf ^fl/fl^* mice at P57+345. [2-way mixed effects ANOVA, axon*genotype interaction: F (1, 5) = 0.2506; p = 0.638; axon F (1, 5) = 111.7; p <0.001; genotype: F (1, 5) = 1.581, p = 0.264]. Graphs show mean ± SD, ^#^p≤0.08, *p<0.05, **p<0.01, ***p<0.001, ****p<0.0001 by Sidak post-test. See also Figure S2

As myelin laid down prior to P57 remains intact in Rα*Myrf ^fl/fl^*mice, acutely preventing adult oligodendrogenesis would not be expected to grossly affect the proportion of callosal axons that are myelinated. However, preventing adult oligodendrogenesis should, over time, lead to Rα*Myrf ^fl/fl^* mice having fewer myelinated callosal axons than control mice. To evaluate this, we collected transmission electron micrographs of the CC from P57+60 or P57+345 control and Rα*Myrf ^fl/fl^* mice (**Fig. 5O-T**). Callosal axon density did not change significantly with age or genotype [P57+60 Control = 2.15 ± 0.26, Rα*Myrf ^fl/fl^* = 1.67 ± 0.63; P57+345 Control = 1.65 ± 0.32, Rα*Myrf ^fl/fl^* = 2.44 ± 0.34 axons/µm^2^ ; 2-way mixed effects ANOVA, age*genotype interaction: F (1, 9) = 9.208; p = 0.014; age F (1, 9) = 0.323; p = 0.58; genotype: F (1, 9) = 0.594, p = 0.46, p>0.1 for all Bonferroni corrected multiple comparisons]. As predicteded, P57+60 control and Rα*Myrf ^fl/fl^* mice had an equivalent proportion of axons that were myelinated (**Fig. 5U**). However, by P57+345, more axons tended to be myelinated in the CC of control than Rα*Myrf ^fl/fl^* mice (**Fig. 5U**; Sidak multiple comparison test, p = 0.0505), but axon diameter was unaffected (**Fig. 5V**).

### Preventing adult oligodendrogenesis does not impair motor performance

We next aimed to determine whether new myelin was required to sustain axon-glial domain integrity or execute established motor behaviours. Unlike genetic approaches that prevent adult oligodendrogenesis by modifying OPC function ^7,61,62^, the conditional deletion of *Myrf* from adult OPCs did not impact nodal integrity, action potential conduction velocity or motor behaviour (**Fig. 6**). When we performed immunohistochemistry to detect NaV1.6^+^ nodes and Caspr^+^ paranodes in the CC (**Fig. 6A-D**), we determined that average node (**Fig. 6E**) and paranode lengths (**Fig. 6F**) and node length distribution (**Fig. 6G, H**) were normal in P57+60 and P57+345 Rα*Myrf ^fl/fl^* mice. Furthermore, the conditional deletion of *Myrf* from adult OPCs was also not associated with a rapid or overt change in the callosal compound action potential (CAP) (**Fig. 6I**). At P57+60, the amplitude (**Fig. 6J**) and half-width (**Fig. 6K**) of the myelinated axon CAP peak was equivalent for control and Rα*Myrf ^fl/fl^*mice, and CAP conduction velocity was unaffected by genotype (**Fig. 6L**).

**Figure 6:**
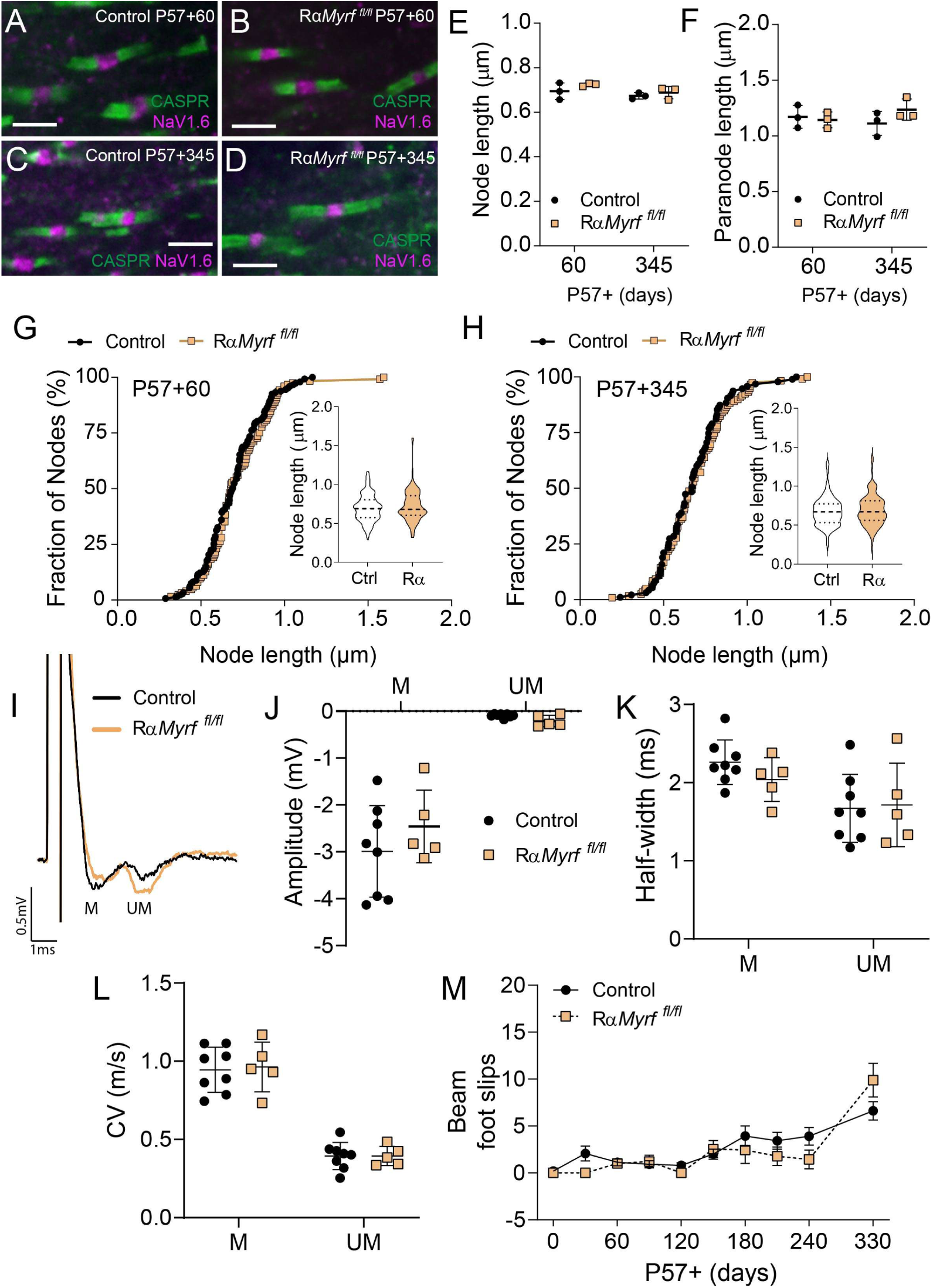
Node and paranode length, and callosal compound action potential conduction is normal in P57+60 RαMyrf ^fl/fl^ mice. (A-D) Single scan confocal image showing nodes of Ranvier (Nav1.6, magenta) and paranodes (caspr, green) in the CC of control and Rα*Myrf ^fl/fl^* mice at P57+60, P57+120 and P57+345. Scale bar represents 2µm (E) Quantification of average node of Ranvier length in the CC of control and Rα*Myrf ^fl/fl^* mice at P57+60 and P57+345. [2-way ANOVA, timepoint*genotype interaction: F (1, 8) = 0.276, p = 0.613; timepoint: F (1, 8) = 3.90, p = 0.083; genotype: F (1, 8) = 2.416, p =0.1587]. Mean ± SD (F) Quantification of average paranode length in the CC of control and Rα*Myrf ^fl/fl^* mice at P57+60 and P57+345. [2-way ANOVA, timepoint*genotype interaction: F (1, 8) = 1.886, p = 0.206; timepoint: F (1, 8) = 0.092, p = 0.76; genotype: F (1, 8) = 0.766, p =0.406]. Mean ± SD (G) Node of Ranvier length distribution in the CC of control (n= 147 nodes) and Rα*Myrf ^fl/fl^* (n=125 nodes) mice at P57+60. Kolmogorov-Smirnov test, K-S D = 0.087, p = 0.68; Mann-whitney U test (inset), M-W U = 8596, p = 0.36. Median (solid line) ± interquartile range (dashed lines) (H) Node of Ranvier length distribution in the CC of control (n=95 nodes) and Rα*Myrf ^fl/fl^* (n=119 nodes) mice at P57+345. Kolmogorov-Smirnov test, K-S D = 0.087, p = 0.81; Mann-whitney U test (inset), M-W U = 5370, p = 0.53. Median (solid line) ± interquartile range (dashed lines) (I) Example callosal CAP traces from control and Rα*Myrf ^fl/fl^* mice at P57+60. (J) Amplitude of the myelinated (M) and unmyelinated (UM) peaks of the callosal CAPS from control and Rα*Myrf ^fl/fl^* mice at P57+60. [2-way ANOVA, axon*genotype interaction: F (1, 22) = 1.557, p = 0.225; axon population: F (1, 22) = 97.98, p <0.0001; genotype: F (1, 22) = 0.634, p =0.43]. Mean ± SD (K) Half-width of the myelinated (M) and unmyelinated (UM) peaks of the callosal CAPS from control and Rα*Myrf ^fl/fl^* mice at P57+60. [2-way ANOVA, axon*genotype interaction: F (1, 22) = 0.035, p = 0.85; axon population: F (1, 22) = 134.8, p <0.0001; genotype: F (1, 22) = 0.035, p =0.85]. Mean ± SD (L) Velocity of the myelinated (M) and unmyelinated (UM) axons in the callosal CAP of control and Rα*Myrf ^fl/fl^* mice at P57+60. [2-way ANOVA, axon*genotype interaction: F (1, 22) = 0.709, p = 0.408; axon population: F (1, 22) = 8.452, p = 0.008; genotype: F (1, 22) = 0.322, p =0.57]. Mean ± SD (M) Quantification of foot slips made by control and Rα*Myrf ^fl/fl^* mice performing the beam walk test at P57-1 to P57+330. [Restricted Maximum Likelihood (REML) mixed effects model: F (1, 22) = 0.749, p = 0.396; age F (3.776, 79.71) =19.76, p < 0.0001; interaction F (9, 190) = 2.445, p = 0.012]. Mean ± SEM. See also Figure S3.

To determine whether adult-born OLs and myelination influence movement, balance or motor coordination, we compared the performance of control and Rα*Myrf ^fl/fl^* mice across a battery of motor function tests, mapping their performance from P57-1, one day before tamoxifen delivery, until P57+300 or P57+330. Consistent with a previous report ^7^, latency to fall from the Rotarod was equivalent for control and Rα*Myrf ^fl/fl^* mice at all timepoints examined (**Fig. S3**). Grid-walk task performance was equivalent for control and Rα*Myrf ^fl/fl^*mice (**Fig. S3**). In the beam-crossing task, control and Rα*Myrf ^fl/fl^* mice experienced a similar age-related increase in foot slips, but performance was not impacted by genotype (**Fig. 6M**). From P57-1 to P57+330, Digigait stride parameters were largely equivalent between control and Rα*Myrf ^fl/fl^*mice (**Fig. S3**), and Rα*Myrf ^fl/fl^* mice did not experience muscle weakness, as their grip strength measures for fore paws or all paws were equivalent to control mice (**Fig. S3**). At P57+60, Rα*Myrf ^fl/fl^* performance in the radial arm maze was also normal (**Fig. S3**). These data suggest that new myelin is not required for mice to continue to perform basic motor tasks and learn in adulthood.

### Adult oligodendrogenesis promotes spine stabilisation on the basal dendrites of layer V pyramidal neurons

To determine if new OLs influence synaptic plasticity in adulthood, even under standard housing conditions, we crossed the Rα*Myrf ^fl/fl^* with *Thy1-YFPH* transgenic mice. This allowed us to visualise and reconstruct the apical (**Fig. 7A**) and basal (**Fig. 7B**) dendrites of layer V pyramidal neurons in P57+60 control and Rα*Myrf ^fl/fl^* mice. At P57+60, spine density along apical and basal dendrites was unaffected by genotype (**Fig. 7C**). Furthermore, along apical dendrites, the density of thin, mushroom and stubby spines (**Fig. 7D**), and the proportion of each spine type (**Fig. 7E**), were unchanged between control and Rα*Myrf ^fl/fl^* mice. By contrast, we found that the density of thin spines was significantly elevated along the basal dendrites of P57+60 Rα*Myrf ^fl/fl^* mice (**Fig. 7F**). As spine density was normal, we confirmed that this changed the ratio of thin to mushroom spines. P57+60 *RαMyrf ^fl/fl^* mice had a significantly higher proportion of thin spines, and a significantly reduced proportion of mushroom spines, compared to age-matched controls (**Fig 7G**).

**Figure 7:**
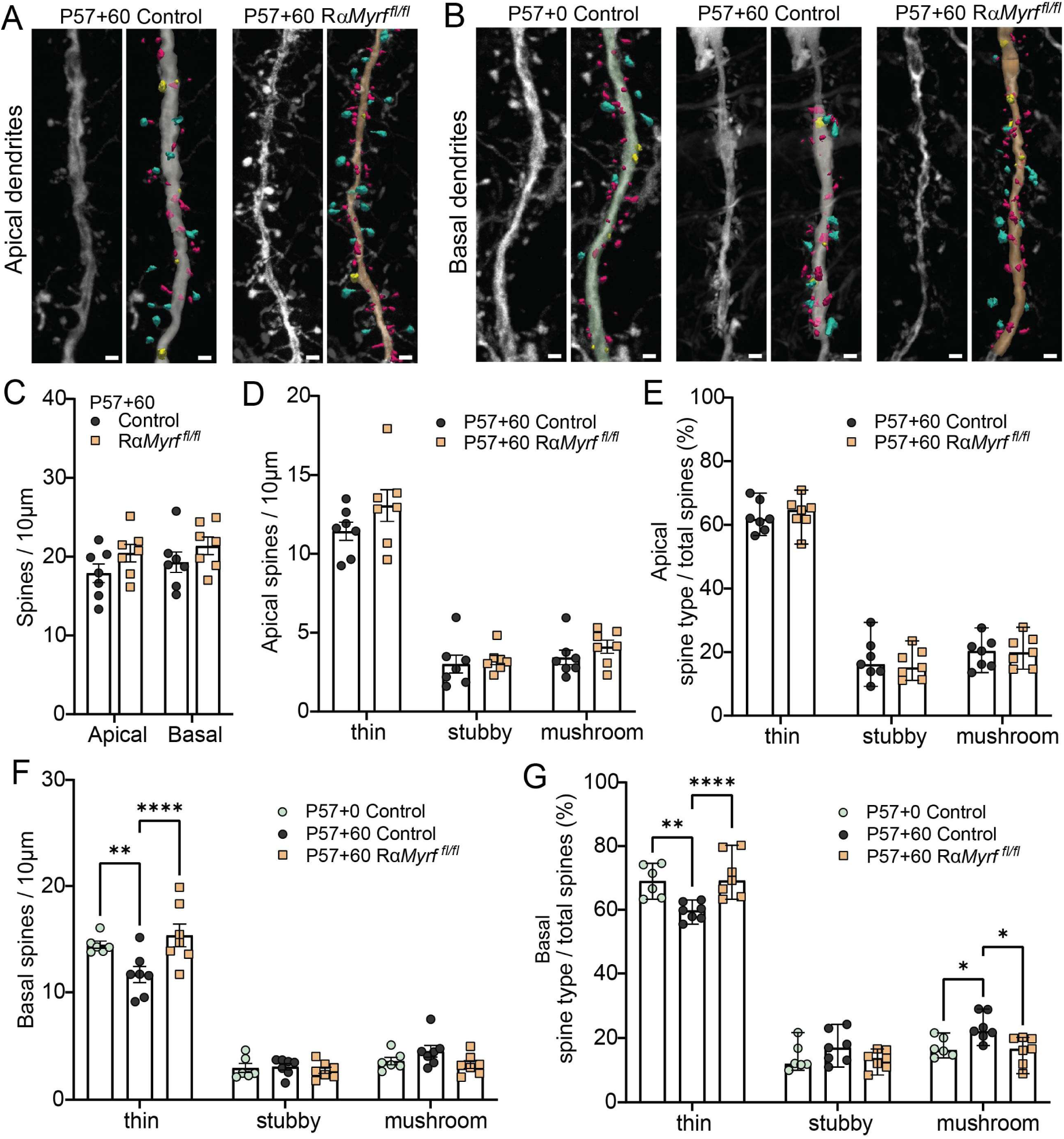
Between 2 and 4 months of age, adult oligodendrogenesis is required for layer V pyramidal neurons to reduce the density of plastic thin spines along basal dendrites. (A) Representative Thy1YFP+ layer 5 pyramidal neuron apical dendrite images (left panels) and rendered Neurolucida-360 traced segments (right panels) from the motor cortex of P57+60 control and P57+60 Rα*Myrf^fl/fl^* . Spine subtype classifications = thin (magenta); stubby (yellow); mushroom (cyan). (B) Representative Thy1YFP+ layer 5 pyramidal neuron basal dendrite images (left panels) and rendered Neurolucida-360 traced segments (right panels) of segments from the motor cortex of P57+0 controls [age-matched P57+60], P57+60 controls and P57+60 RαMyrf^fl/fl^ mice. (C) Mean spine densities (number spines per 10µm dendrite length) for apical and basal dendrites in P57+60 control and P57+60 Rα*Myrf^fl/fl^* mice. [2-way mixed ANOVA: dendrite*group interaction F(1, 12) = 0.055 p=0.81, dendrite F(1, 12) = 1.372 p = 0.26, group F(1, 12) = 3.043 p = 0.107] (D) Mean densities for each spine type (thin, mushroom, stubby) per 10µm apical dendrite length in P57+60 control and P57+60 Rα*Myrf^fl/fl^* mice. [2-way mixed ANOVA: spine type*group interaction F(2, 24) = 0.799 p=0.46, spine type F(2, 24) = 166.1 p <0.001, group F(1, 12) =2.537 p = 0.137] (error bars = SEM). (E) Median percent of spines in each morphological classification (thin, mushroom, stubby) per apical dendrite (95%CIs). The percent of spines in each classification per analysed apical dendrite segment was not significantly different between genotypes [MWU tests to compare medians, P57+60 control vs. P57+60 Rα*Myrf^fl/fl^* in each type, all Bonferroni corrected p-values >0.05]. (F) Mean basal dendritic spine densities of each morphological spine type classification (thin, mushroom, stubby) in the motor cortex of P57+ 0 controls (green circles); P57+60 control (black circles), P57+60 RαMyrf^fl/fl^ (orange squares) [2-way mixed ANOVA with group (between-subjects), and spine type (within-subjects: thin, mushroom, stubby); type*group interaction: F(4,34) = 6.384, p =0.001; group: F(2,17) = 1.209, p = 0.323; type: F(2,34) = 360.02, p < 0.001]. (error bars = SEM). (G) Median proportion of spine type (thin, mushroom, stubby) per basal dendrite (95%CIs) from motor cortex of P57+0 controls (green circles); P57+60 control (black circles), P57+60 RαMyrf^fl/fl^ (orange squares) mice. [thin spines KW: *H*(2,20) = 13.08, p = 0.001; P57+60 control vs P57+60 Rα*Myrf^fl/fl^* mice (z = -3.30, Bonferroni-adjusted p = 0.003) and P57+0 control vs P57+60 Rα*Myrf^fl/fl^* mice (z = -2.89, Bonferroni-adjusted p = 0.012); mushroom spines [KW: *H*(2,20) = 9.81, p = 0.007], with P57+0 controls vs P57+60 controls (z = 2.58, p = 0.03), P57+60 control vs P57+60 Rα*Myrf^fl/fl^* (z = 2.8, p = 0.015), P57+0 vs P57+60 Rα*Myrf^fl/fl^* (all p-values >0.99)] The percentage of stubby type spines did not differ significantly between groups (p>0.05). Datapoints represent individual subjects (mean of ≥ 5 dendrite segments per mouse). Scale bars = 1μm. Asterisks denote Bonferroni-adjusted p-values: *p<0.05, **p<0.01, ***p<0.001.

To determine whether this is because a proportion of thin spines normally transition and accumulate as mushroom spines over time in control mice, and this process fails in Rα*Myrf ^fl/fl^* mice, or mushroom spines are lost over time in Rα*Myrf ^fl/fl^* mice, we imaged and reconstructed the basal dendrites of P57+0 *Thy1-YFPH* control mice. We report that basal dendritic spine density did not change with time [One-way ANOVA F(2,17) = 1.14, p = 0.34; P57+0 control mice had 21.02 ± 1.2 spines/10µm; P57+60 control mice had 19.28 ± 3.41 spines/10µm; P57+60 Rα*Myrf ^fl/fl^*mice had 21.36 +/- 2.93 spines/10µm]. However, P60+0 control basal dendrites had a higher density of thin spines than the P57+60 controls, resembling the composition of P57+60 Rα*Myrf ^fl/fl^* basal dendrites (**Fig. 7F**). Conversely, a lower proportion of P57+0 control basal dendritic spines were stable mushroom spines, relative to P57+60 controls, but the proportion was equivalent to that of P57+60 Rα*Myrf ^fl/fl^* mice (**Fig. 7G**). These data indicate that new myelin addition is required for layer V MC neurons to increase their proportion of stable mushroom spines across adulthood.

## Discussion

The addition of new myelin to the adult brain supports the acquisition of motor skills ^7,10^ and long-term memory recall ^33,34^. However, it is unclear whether the ability of mature OLs to maintain myelin or the addition of new myelin impacts the strength and stability of synaptic connections. By deleting *Myrf* from mature OLs at P57, we disrupted the myelin maintenance program in OLs generated before young adulthood. Over the following 60 days, myelin basic protein levels were reduced, CAP conduction slowed (**Fig. 1**), and mice developed fine and gross motor dysfunction that impaired their ability to execute basic as well as established, complex movement (**Fig. 2**). This was accompanied by changes in the postsynaptic structure of layer V MC pyramidal neurons, and their response to glutamate. Apical or basal dendritic spine density was unchanged, but each dendritic compartment had an increased proportion of stable, mushroom spines (**Fig. 3**), and mEPSC amplitude was increased (**Fig. 4**). The significant skewing of spine morphology towards mushroom spines, suggests that the OL myelin maintenance program is critical for neuron homeostasis and their ability to support a majority of highly plastic spines to remain adaptable throughout life. Conversely, when *Myrf* was instead deleted from adult OPCs, to prevent new OL maturation and further myelin addition (**Fig. 5**), CAP conduction velocity was normal, and the mice were still able to execute basic as well as established, complex motor tasks (**Fig. 6**). However, 2-months without adult myelination was sufficient to alter the structure of layer V pyramidal neurons in the MC. In particular, the basal dendrites of P57+60 Rα*Myrf ^fl/fl^* retained the higher proportion of thin, transient spines, that was characteristic of P57 but not P117 control dendrites, suggesting that the basal dendritic spine configuration was “frozen” at the time of *Myrf* deletion. These data indicate that adult myelination supports the potentiation and stabilisation of basal dendritic spines over time, consistent with a role in consolidation.

### OL myelin maintenance allows layer V pyramidal neurons to balance synaptic plasticity and stability

At P57+60, the apical and basal dendrites of PLP*Myrf ^fl/fl^*mice had an elevated proportion of large mushroom spines (**Fig. 3**). As the size of the postsynaptic spine head predicts synapse stability, with larger spines having a reduced likelihood of elimination ^56,63^, these data suggest that more of the PLP*Myrf ^fl/fl^* layer V pyramidal dendritic spines are stable. The large mushroom dendritic spines also have the highest concentration of excitatory AMPA receptors at the postsynaptic density ^55^, which is supported by an increase in mEPSC amplitude in *PLPMyrf ^fl/fl^*layer V neurons (**Fig. 4**). This increased responsiveness to glutatmate may explain previous reports that myelin loss is associated with an increase in cfos expression by excitatory neurons in MC ^64^, and an increase in sEPSC generation in the striatal medium spiny neurons ^65^, which are the downstream of MC layer V neurons ^41^.

A larger number of stable mushroom spines could reflect altered Hebbian or homeostatic plasticity ^66^. For example, an increase in the activity of specific excitatory inputs could strengthen those spines. Alternatively, if the conditional deletion of *Myrf* from OLs produced a sustained decrease in network activity, the reduced layer V pyramidal neuron firing would trigger a homeostatic response that would increase AMPA receptor expression in spines (increasing spine size) to restore neuronal firing ^66^. It is also possible that preventing myelin maintenance could change the expression of neuromodulators like norepinephrine or dopamine, which can change the threshold for long-term-potentiation ^67,68^. Irrespective of the cause, the conditional deletion of *Myrf* from OLs produced a major structural and functional change to the MC output neurons, that would render them sensitive to changes in glutamate expression or signalling. Such changes could contribute to neurodegeneration in demyelinating diseases like multiple sclerosis (MS) ^69^.

While dendritic spine morphology was significantly altered in PLP*Myrf ^fl/fl^* mice, spine density was preserved (**Fig. 3**). By contrast, synapse loss has been reported in the brain, spinal cord and retina of people with MS ^70^ ^71–73^ prior to neuron loss ^72,74,75^. Synapse loss has also been reported in rodent models of inflammatory demyelination ^75^, however, the outcome is highly variable with some neurons maintaining synapse density and others even increasing synapse density ^76,77^. As demyelination does not appear to drive synapse loss directly ^75,78,79^, and our data suggest that myelin loss would reduce rather than increase the likelihood of spine removal ^63^, the outcome will likely depend on the inflammatory environment. In a demyelinating disease context, increased microglial synapse engulfment would be enhanced by blood brain barrier dysfunction and elevated complement C3 expression Werneburg, 2020 #521;Merlini, 2019 #522}.

### Adult myelination supports synapse stabilisation and reduces plasticity

Preventing OL addition between P57 and P117 changed the morphological composition of spines along the basal dendrites of layer V pyramidal neurons in MC. More specifically, P57+60 Rα*Myrf ^fl/fl^* mice had dendrites that resembled P57 control rather than P57+60 control mice, as they retained a higher proportion of immature, transient, thin spines (**Fig. 7**). This is reminiscent of a previous report showing that without myelin addition from development (P10) through to adulthood, pyramidal neurons in the visual cortex retained their adolescent-level of plasticity ^25^. Spine structural plasticity is heavily influenced by developmental stage, being less restricted in juvenile than adult mice ^56,80–82^. This is reflected by an increased proportion of spines having a stable mushroom morphology with age. Live imaging of the apical tufts of layer II/III and V dendrites in the mouse somatosensory cortex revealed that the proportion of stable spines increases from ∼35% at P25, to ∼54% at P80, ∼66% at P120 and ∼73% at P225 ^56^. Our data indicate that adult myelination promotes spine stabilisation over the lifespan but also shows that it exerts a greater influence over spine stabilisation in the basal dendritic compartment.

Layer V MC pyramidal neurons receive monosynaptic inputs from layers I, II/II and V M1 neurons, as well as neurons in the frontal association, motor and somatosensory cortices, thalamus, basal ganglia and other subcortical regions ^38,83^. From each region, the neurons that synapse with the apical or basal dendrites of layer V MC neurons are almost discrete populations ^38^. The stabilisation of new spines on apical dendrites corelates with motor task performance and their artificial elimination impairs performance of the learned motor task ^36^. However, the specific role of basal inputs to layer V MC neurons during task acquisition versus recall requires further investigation. A recent study found that spines along the apical and basal dendrites of layer II/III MC pyramidal neurons applied distinct activity-dependent plasticity rules ^54^. The strengthening of apical synapses was predicted by their coactivity with other local synapses, while the strengthening of basal dendrites was predicted by their activity being coincident with the postsynaptic action potential ^54,84^. If layer V MC neurons apply similar compartment-specific functional synaptic organization, our data could indicate that adult myelination increases the likelihood that basal synapse activity will be coincident with the postsynaptic action potential, to support the transition of select inputs from transient thin to stable mushroom spines over time. While this is not a learning study, synaptic plasticity on layer V pyramidal neuron dendrites plays a crucial role in the acquisition of new motor skills ^37,42,85,86^, and the modulation of synaptic plasticity in discrete dendritic compartments may explain the ability of adult myelination to support learning and memory functions ^7,33,34^.

## Acknowledgements

We thank Mr Mackenzie Clutterbuck and the University of Tasmania Animal Services team for assistance with animal care and genotyping. This research was supported by grant and fellowship funding from: the National Health and Medical Research Council (1030939, 1045240, 1077792 and 2012140); the Australian Research Council (DP180101494; DP220100100); the Medical Research Future Fund (EPCD08); Multiple Sclerosis Australia (11-014, 15-054, 16-105, 17-0223, 19-0696, 20-460; 21-4-017); the Menzies Institute for Medical Research, University of Tasmania, and the Mater Foundation, Equity Trustees and the Trusts of L G McCallum Est. The funding sources had no role in the study design; collection, analysis or interpretation of the data; in writing the manuscript or in the publication process.

## Author contributions

KMY, CLC, BE and REP conceived the project. KMY, CLC, BE and KM acquired funding. BE and CB provided resources. CB, REP, CLC, KMY and KM developed methodology. REP, KM, KAP, RR, PTN, CLC and KMY carried out the experimental investigation. REP, KM and CLC were involved in data curation. REP, KM, CLC, AN and KMY conducted formal analysis and visualized the data. KMY undertook project administration. KMY and CLC provided supervision and wrote the manuscript.

## Declarations of interest

The authors declare no competing interests.

## STAR Methods

### Key resources table

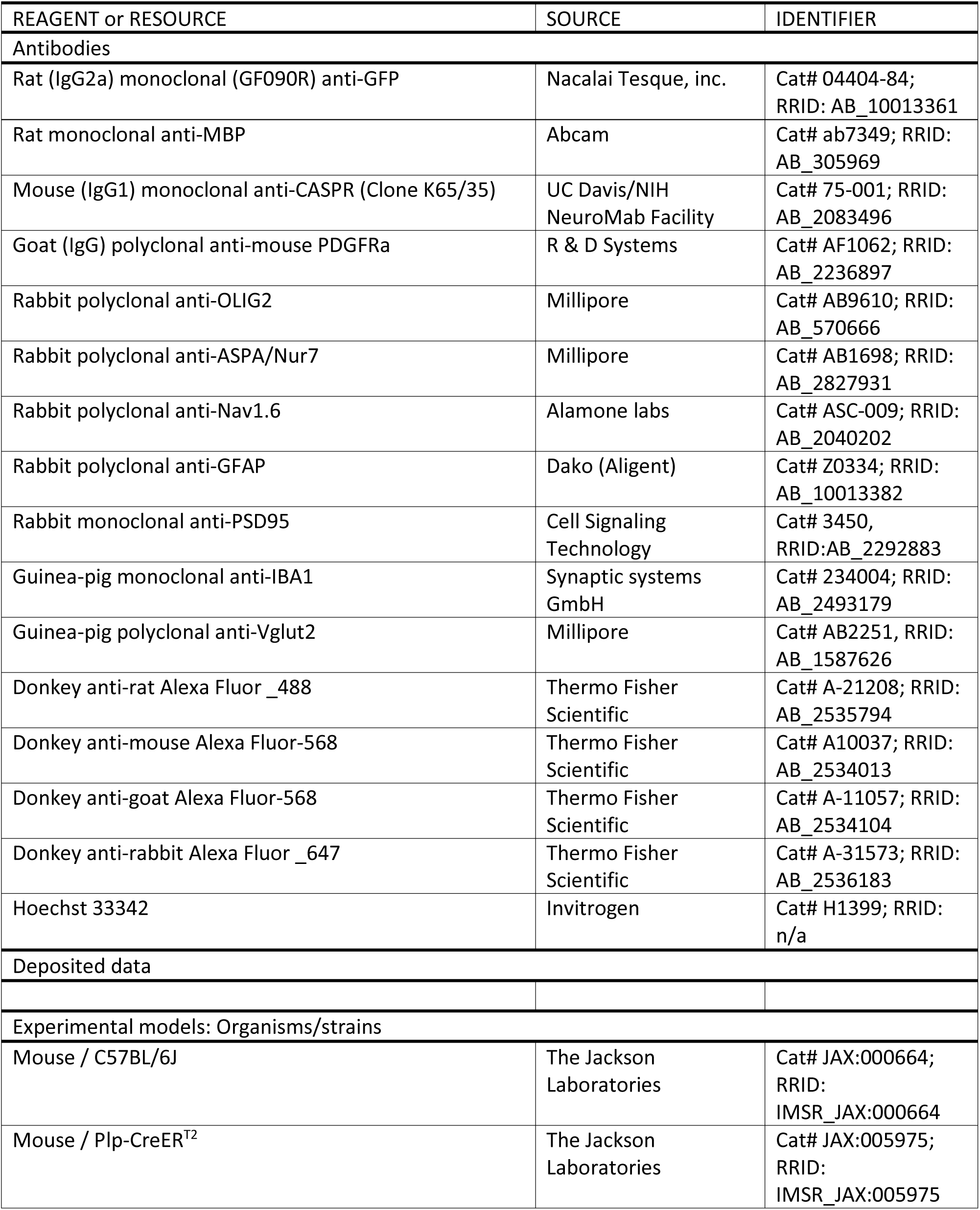
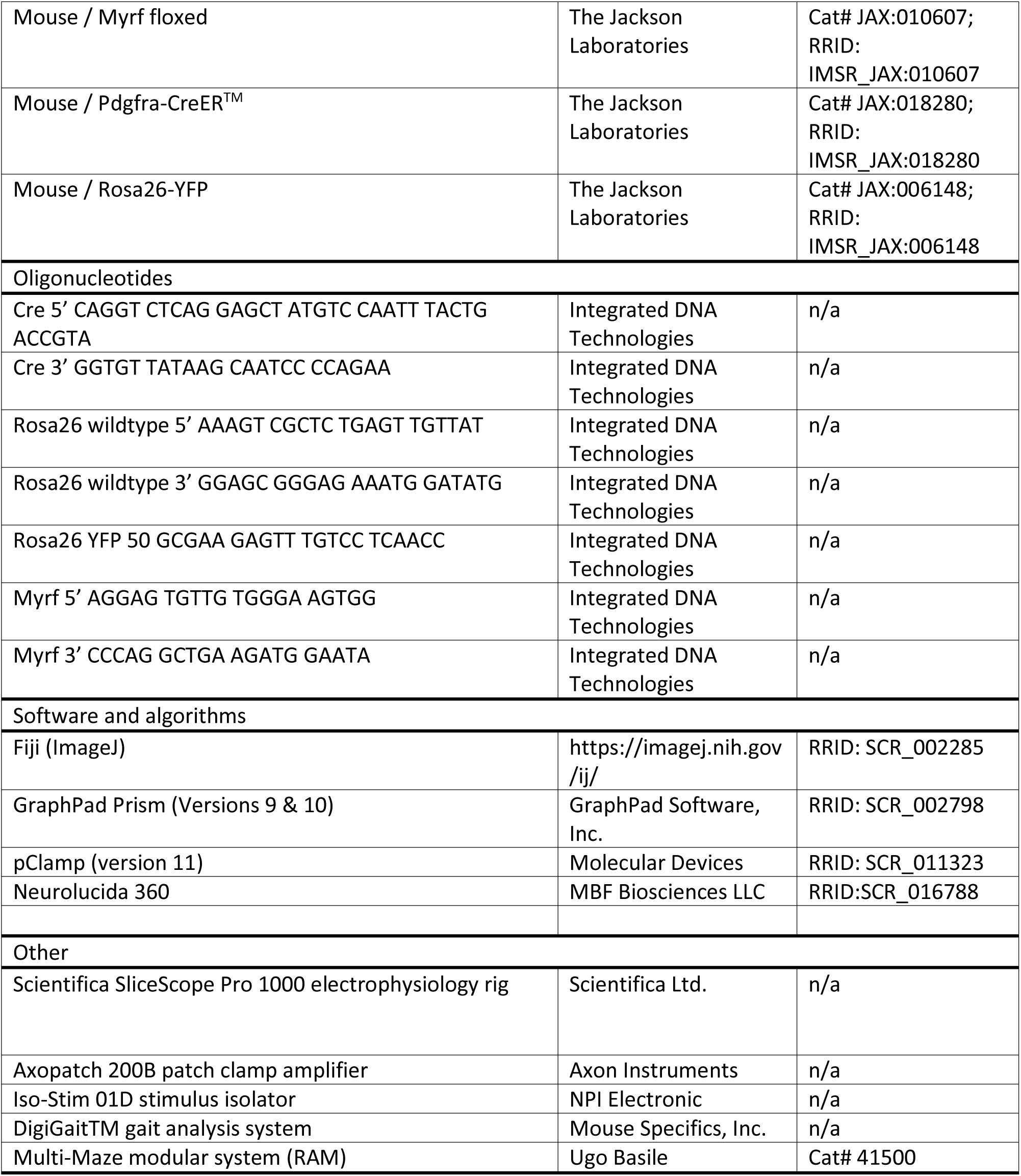

### Resource Availability

#### Lead contact

Further information and requests for resources and reagents should be directed to and will be fulfilled by the lead contact, Kaylene Young.

#### Materials Availability

This study did not generate new unique reagents

#### Data and code availability

All data generated and analysed for this study are included in the manuscript. Raw image files are available upon reasonable request from the corresponding author.

### Experimental Model and Subject Details

All animal experiments were approved by the University of Tasmania Animal Ethics Committee (A0013471 and A0016151) and were carried out in accordance with the Australian code of practice for the care and use of animals for scientific purposes. *Plp-CreER^T2^* ^87^, *Pdgfrα-CreER^TM^* ^32^ (JAX:018280), *Rosa26-YFP* ^88^ (JAX:018280; RRID: IMSR_JAX:018280) and *Thy1-YFPH* ^89^ (JAX:003782) transgenic mice were purchased from Jackson Laboratories and *Myrf ^fl/fl^* transgenic mice ^46^ were provided by Ben Emery (Oregon Health and Science University). Each was maintained as a live colony on the C57BL/6 background at the University of Tasmania for >10 years. Mice were interbred to generate experimental mice that were not weaned prior to P35 to ensure normal myelin development. Mice were group housed (2-5 per cage) in same sex groups in individually ventilated Optimice^TM^ cages on a 12-hour light/dark cycle with free access to food and water. Mice bred for behavioural experiments were handled for ∼5 min daily for two weeks prior to commencing the experiment.

### Method Detail

#### Genomic DNA extraction and PCR genotyping

Genomic DNA was extracted from mouse tissue biopsies and 50-100ng of the genomic DNA used to perform polymerase chain reaction (PCR) to detect the relevant transgenes as previously described ^90^. The following primers were used to detect the *Rosa26-YFP* transgene: Rosa26 wildtype 5’ AAAGT CGCTC TGAGT TGTTAT; Rosa26 wildtype 3’ GGAGC GGGAG AAATG GATATG and Rosa26 YFP 5’ GCGAAGAGTTTGTCCTCAACC [94°C for 4’; 37 cycles of 94°C for 30”, 60°C for 45” and 72°C for 60”, and a final 72°C for 10 min]. *Pdgfrα-CreER^TM^* and *Plp-CreER^T2^* mice were genotyped using primers that bind to *Cre recombinase*: Cre 5’ CAGGT CTCAG GAGCT ATGTC CAATT TACTG ACCGTA and Cre 3’ GGTGT TATAA GCAAT CCCCA GAA [94°C for 4’, 34 cycles of 94°C for 30”, 62°C for 45” and 72°C for 60”, and a final 10 min at 72°C]. The *Myrf floxed* gene was detected with Myrf 5’ AGGAG TGTTG TGGGA AGTGG and Myrf 3’ CCCAG GCTGA AGATG GAATA [94°C for 4’, 34 cycles of 94°C for 30”, 62°C for 45” and 72°C for 60”, and a final 10 min at 72°C]. Mice carrying the *Thy1-YFPH* transgene were genotyped by RT-PCR (Transnetyx).

#### Tamoxifen administration and tissue preparation

Tamoxifen (Sigma-Aldrich) was dissolved to 40 mg/ml in corn oil in a sonicating water bath (≥ 1 hour until dissolved). For conditional gene deletion and lineage tracing, postnatal day (P)57 mice received 300mg/kg of Tamoxifen by oral gavage, daily for 4 consecutive days. Mice were analysed at various time-points following Tamoxifen administration, and the number of days is indicated in the text using the nomenclature P57+(days).

For tissue collection, mice were terminally anaesthetised with a 180mg/kg intraperitoneal (i.p.) injection of sodium pentobarbitone and transcardially perfused at ∼9ml/min with 4% paraformaldehyde (PFA; Sigma-Aldrich) (w/v) in phosphate buffered saline (PBS). Brains were removed post-mortem and the 2mm coronal brain slices, generated using a brain matrix (Kent Scientific), were immersion fixed in 4% PFA (w/v) in PBS for 90 min at ∼21°C. Tissue was cryoprotected by immersion in 20% sucrose (Sigma-Aldrich) (w/v) in PBS overnight at 4°C prior to embedding in optimal cutting temperature cryomatrix (Thermo Scientific) and storage at -80°C.

#### Immunohistochemistry, confocal microscopy and quantification

Coronal brain cryosections were generated using a Leica cryostat and processed as floating sections. For immunohistochemistry, we collected 30µm cryosections (20µm for nodal and paranodal analyses). Cryosections were immersed in PBS blocking solution [0.5% (v/v) Triton X-100 and 10% fetal calf serum in PBS] for 1 h, at ∼21°C. Primary antibodies [goat anti-PDGFRα (1:100; GeneTex, California, USA), rabbit anti-ASPA (1:200; Millipore, RRID: AB_2827931); rabbit anti-OLIG2 (1:400 Millipore), rat anti-GFP (1:2000; Nacalai Tesque, Kyoto, Japan), rabbit anti-NaV_1.6_ (1:500 Alomone Labs), and mouse anti-CASPR (Clone K65/35; 1:200 NeuroMab, California, USA)] were diluted in PBS blocking solution and applied to the cryosections overnight at 4°C. Sections were washed thrice in PBS before the application of secondary antibodies conjugated to AlexaFluor-488, -568, -647, -800 (Invitrogen) [donkey anti-goat (1:1000), donkey anti-rabbit (1:1000), donkey anti-mouse (1:1000), and donkey anti-rat (1:500), donkey anti-guinea pig (1:400)] and the nuclear label Hoechst 33342 (1:1000; Invitrogen), diluted in PBS-blocking solution overnight at 4°C. All sections were again washed in PBS before being mounted using DAKO fluoromount.

Immunohistochemistry images were collected using an UltraView Nikon Ti Microscope with Volocity Software (Perkin Elmer), Andor Confocal microscope with Nikon Software (Andor Technology Ltd., Belfast, Northern Ireland), or FluoVIEW slide scanner (Olympus Australia Pty Ltd). Confocal images were collected using standard excitation and emission filters for DAPI, FITC (AlexaFluor-488), TRITC (AlexaFluor-568) and CY5 (AlexaFluor-647), and 20x air objective lens, and for cell quantification, we collected images at 3µm z-intervals that spanned an anatomically defined region of interest. Images were stitched together in the x-y plane to make a single composite image of the region for each brain hemisphere.

Imaging and quantification of node of Ranvier and paranode length was performed as previously described ^49^. Single z-plane Images were collected used a 100x oil immersion lens, and nodes and paranodes were selected for quantification only when a node and its flanking paranodes were intact (flat) within the single z-plane. Length measurements were made using Image J (NIH) while cell number quantification was undertaken using Adobe Photoshop. Cell counts are presented as a density or a percentage of recombined cells. Densities were calculated in mm^2^, where x-y measurements were taken from defined region of interests and z-plane was not included as all tissue for cell quantification was a set tissue thickness (30µm). Percentages were used to represent the proportion of differentiated oligodendrocytes (YFP^+^, PDGFRα-negative) relative to the total number of recombined cells (YFP^+^, OLIG2^+^ cells).

#### Transmission Electron Microscopy

Mice were terminally anaesthetised with a 180mg/kg intraperitoneal (i.p.) injection of sodium pentobarbitone and transcardially perfused at ∼9ml/min with Karnovsky’s fixative (2.5% glutaraldehyde, 2% PFA, 0.25mM CaCl_2_, 0.5mM MgCl_2_ in 0.1M sodium cacodylate buffer). The brains were removed post-mortem and cut into 1mm-thick coronal slices, using a brain matrix (Kent Scientific), that were post-fixed in Karnovsky’s fixative for 2 hours at 21°C. The tissue slices were rinsed and stored in 0.1M sodium cacodylate buffer at 4°C overnight. The corpus callosum was dissected and incubated in 1% osmium tetroxide (OsO_4_) / 1.5% potassium ferricyanide [K_3_Fe(III)(CN)_6_] in 0.1M sodium cacodylate buffer in the dark for 2 hours at 4°C, before being dehydrated in ethanol and propylene oxide, and embedded in Epon812 resin. 70nm ultrathin sections, transecting the corpus callosum, were cut using a Leica Ultra-cut UCT7, and stained with uranyl acetate and lead citrate for imaging on a Hitachi HT7700 transmission electron microscope. G-ratio, the proportion of myelinated axons, mitochondrial density and myelin degeneration were quantified from images collected at 8kV. Image analysis was carried out using Image J (NIH) and involved measuring ≥ 100 axons per mouse. The number of myelin wraps was quantified from images collected at 80kV and involved counting the major dense lines for ≥ 21 transected myelinated axons per mouse.

#### Motor behaviour

Motor behaviour was evaluated as previously described ^62^. Briefly, male and female mice of all genotypes were first evaluated prior to tamoxifen administration (P57-1) to determine baseline performance. Repeat motor testing was performed until P57+345. Mice were tested at the same time of day (14:00-17:00), during the light phase of the 12-hour light/dark cycle, and in low-light conditions (∼20 lux). Mice were habituated to the testing room for at least 1 h before commencing.

##### Rotarod

Balance and coordination was assessed using the RotaRod Advanced apparatus (TSE Systems). Mice were trained for two weeks at rotation speeds of 10, 20 and 40rpm, with an acceleration of 8 rpm/minute prior to first data collection to accommodate for the learning phase of the task. Following habituation, mice were tested using a RotaRod paradigm ^50^ in which the rotating rod increased speed from 4 to 40rpm over 300 s, followed by a further 60 s at 40 rpm. For all experiments ‘latency to fall’ (time a mouse falls off the rotating rod) was recorded. Rotarod performance testing was repeated three times per session with a 15-min inter-trial-interval.

##### Grid walk

Mice were placed onto a horizontal, 2 cm wire mesh grid elevated 100 cm above the ground and were left to explore for 5 minutes. Each mouse was video recorded from underneath the grid using a GOPRO^TM^ Hero5 and the number of missteps (the stepping foot slips through the grid) quantified in videos by an experimenter blind to genotype. We calculated the total number of missteps within the first 100 steps, or over the 5 minutes if 100 steps were not taken, and this was expressed as a percentage of steps analysed.

##### Beam walk

Mice were trained to cross a 100 cm length of wooden dowel (10 mm diameter) that was fixed horizontally and elevated 50 cm above the surface of the bench. Individual trials were video recorded at 60 frames/s using a GOPRO^TM^ Hero5 handheld camera. The videos were then scored by an experimenter blind to genotype. The total number of foot slips that occurred in a single crossing was recorded for each mouse. A foot slip was defined as any paw falling below the level of the beam.

##### Gait analysis

Running gait was assessed using the DigiGait^TM^ Imaging System (Mouse Specifics, Inc., MA, USA). Briefly, mice were individually evaluated, by placement within the brightly lit plexiglass chamber on the transparent motorized treadmill belt and allowed 5 min to habituate to the chamber. The treadmill was then turned on at a speed of 5cm/s for 5 seconds before the belt speed slowly increased to 22 cm/s over a 10 second period. A 30 second video of the mouse running at 22 cm/s was recorded at 164 frames/s using a Basler scA640-74fc digital camera mounted underneath the treadmill. Analysis of 3-4 second video segments (∼10 steps) were analysed using DigiGait^TM^ Analyzer software version 15 (Mouse Specifics, Inc., MA, USA). Data for the left and right limbs were pooled, but forelimbs and hindlimbs were analysed separately.

##### Grip strength

A grip strength meter (Melquest, cat. #GPM-100) was used to estimate muscular strength as previously described ^91^. Each mouse was weighed, positioned ready to grasp the grid mounted on the force gauge, and the equipment reset to 0 gF after stabilization. Each mouse was allowed to grasp the grid with all four paws, pulled back over the grid, and the maximum tension recorded. A test was repeated if all four paws failed to successfully grasp the grid or the mouse bit or knocked the grid. Five successful trials were recorded per mouse. We calculated force (gF) relative to body weight (g) for each mouse by dividing the mean of five successful tests by body weight.

### Dendritic spine imaging and quantification

Mice carrying the *Thy1-YFPH* transgene were transcardially perfused as described above, with the brain slices light-protected during post-fixation and cryoprotection. Coronal cryosections (30 µm) containing the MC were thaw-mounted directly onto slides and allowed to dry for 15 minutes before submersion in PBS for 3 minutes to remove cryomatrix, then briefly submerged in dH_2_O to remove salts. Sections were then allowed to dry before coverslipping with Prolong Glass Antifade mounting medium (Thermo Fisher Scientific) and high precision coverslips (No.1.5H, Marienfeld Superior). For blinding, slides were re-labelled prior to imaging.

Dendrite segments were imaged on an Olympus FLUOVIEW FV3000 (BX63LF) laser scanning microscope (514nm excitation), with high-numerical aperture oil-immersion objective (Olympus Super Apochromat UPLSAPO-100XO/1.4). Thy1-YFP^+^ dendrites in the MC were selected pseudo-randomly, with the criteria that the dendrite segment was relatively flat in the x-y plane, and sufficiently isolated from other YFP^+^ soma and processes that individual spines could be identified. Apical dendrite segments were within layers 5 and 2/3. Basal dendrite segments were selected only when the soma of origin in layer 5 could be identified and captured in the same field of view. Images were 1024×1024 pixels, at 0.07- 0.1µm/pixel, and z-step size of 0.09 to 0.2µm, acquired at 0.2µs/pixel dwell time and averaged 3x by frame. Because YFP^+^ signal varied, laser power, detector voltage and offset parameters were adjusted for each image to avoid saturated pixels and minimise background.

Dendritic spine reconstructions were performed in Neurolucida 360 (v.2021.1.3, MBF Bioscience LLC, Williston, VT USA). Traced dendrite segments were at least 10µm in length, and at least 5 dendrites were reconstructed per animal. Dendrites were reconstructed with the ‘user-guided’ manual tree tool then adjusted to accurately fill dendrite volumes. Individual spines were manually selected, adjusting detector sensitivity for each spine to ensure the spine head was accurately reconstructed, with other detection parameters set at 10 voxel minimum size, 0.3µm minimum height and 2.5µm maximum distance from dendrite. Spines were semi-automatically classified into morphological subtypes of thin, mushroom or stubby, using the Neurolucida 360 multi-dimensional classifier, with settings of head-to-neck ratio of 1.1, length-to-head ratio of 2.2-2.3, and mushroom head size >0.3µm. ∼15% of spines could not be automatically classified accurately, usually due to patchy fluorescence in the spine neck, and were manually classified based on measured head diameter across the widest portion, and head-neck appearance. Dendritic spine densities and proportions of spines classified as each subtype were obtained by batch analysis in Neurolucida Explorer (v.2020.3.1) calculated for each traced dendrite, then averaged per animal (≥ 5 dendrite segments per mouse). Image processing and analysis was performed by a researcher blind to experimental group.

### Electrophysiology

#### Extracellular CAP recordings

CAPS were recorded from the adult mouse corpus callosum as previously described ^49^. Briefly, PLP*Myrf ^fl/fl^* and Rα*Myrf ^fl/fl^* mice were killed by cervical dislocation 35-42 days (P57+35) or 60-67 days (P57+60) after Tamoxifen administration, respectively. Brains were rapidly dissected into ice-cold sucrose solution containing in mM: sucrose 75, NaCl 87, KCl 2.5, NaH_2_PO_4_ 1.25, NaHCO_3_ 25, MgCl2 7, and CaCl_2_ 0.95. Live coronal brain vibratome (Leica VT1200s) slices (400µm) were generated spanning Bregma +0.8 and −0.2. Slices were incubated at ∼32°C for 45 min in artificial cerebral spinal fluid (ACSF) containing in mM: NaCl 119, KCl 1.6, NaH2PO_4_ 1, NaHCO_3_ 26.2, MgCl_2_ 1.4, CaCl_2_ 2.4, and glucose 11 (300 ± 5 mOsm / kg), then transferred to ∼21°C ACSF saturated with 95% O_2_/5% CO_2_.

CAPs were evoked by constant current, stimulus-isolated, square wave pulses (200 ms duration, delivered at 0.2 Hz), using a tungsten bipolar matrix stimulating electrode (FHC; MX21AEW), and detected using glass recording electrodes (1-3 MΩ) filled with 3M NaCl. To quantify CAP amplitude, the asymptotic maximum for the short-latency negative peak (myelinated peak, M) was determined with stimulating and recording electrodes placed 1mm apart and varying the stimulus intensity (0.3–4.0 mA) using an external stimulus isolator (ISO-STIM 1D) before recording at 80% maximum stimulation. To enhance the signal-to-noise ratio, all quantitative electrophysiological analyses were conducted on waveforms that were the average of eight successive sweeps, amplified, and filtered (10 kHz low pass bessel) using an Axopatch 200B amplifier (Molecular Devices), digitized at 100 kHz and stored on disk for offline analysis.

The CV of myelinated (M) and unmyelinated (UM) axons was estimated by changing the distance between the stimulating and recording electrodes from 1 to 3 mm spanning the CC, while holding the stimulus intensity constant (80% maximum). The peak latency of the M and UM axons was measured at each point and graphed relative to the distance separating the electrodes. A linear regression analysis was performed to yield a slope, inverse to the velocity for each brain slice. The average velocity for both CAP components (M, UM) was determined for each animal and this value was used for statistical comparison.

#### Whole cell patch clamp electrophysiology

Acute brain slices were generated from PLP*Myrf ^fl/fl^* mice 65-71 days after tamoxifen delivery (P57+65), and mini excitatory post-synaptic currents (mEPSCs) recorded from layer V pyramidal neurons in the adult mouse MC as previously described ^92^. Briefly, animals were killed by cervical dislocation, brain rapidly dissected into ice cold sucrose cutting solution (as described above) bubbled with carbogen. 300µm coronal brain slices, containing the motor cortex, were cut on a vibratome and transferred to 37°C ACSF (in mM: NaCl 126, KCl 1.6, NaH_2_PO4 1.1, MgCl2 1.4, NaHCO3 26, CaCl2 2.4, glucose 11) for 30 minutes, then at ∼21°C for at least 1hr. Whole cell patch clamp recordings were performed at ∼21°C, using a 4-6 MΩ micropipette glass electrode filled with caesium internal solution (in mM: CeMeSO_4_ 125, NaCl 4, KCl 3, MgCl 6H_2_0 1, HEPES 8, EGTA 9, phosphocreatine 10, MgATP 5, Na_2_GTP 1), and external solution of ACSF with 3mM TTX (Abcam) and 1mM picrotoxin. Data were recorded using an Axon Instruments amplifier, sampled at 50kHz and high pass filtered at 1kHz. Layer V pyramidal neurons within the MC were identified by morphology and membrane resistance between 120-150MΩ, and membrane capacitance >70pF. Cells were excluded if: access resistance (Ra) was >20MΩ or changed by more than 15% during recordings; or if the response and return to baseline for a hyperpolarising test pulse (−500pA) was unstable 3 minutes after sealing. Data were analysed using Axograph X (version 1.8.0) to automatically detect mEPSCs within 2.5 standard deviations of a pre-defined variable-amplitude template and to measure the amplitude of detected events. Amplitude was measured relative to baseline (mean of 6ms immediately before each event). Events with negative or zero decay times or decay times above 100 ms were removed post-processing.

### Statistical analyses

The number of mice analysed in each group (*n*) or the number of cells, axons, nodes or paranodes is indicated in the corresponding figure legends. All statistical analyses were performed using Prism 9 (GraphPad Software) or SPSS v.22 (IBM). Data were further analysed by parametric or nonparametric tests as appropriate. Data comparing two groups at a single time point were analysed using a parametric two-tailed t-test or a non-parametric Mann-Whitney U (MWU) test. Data distributions were compared using a Kolmogorov–Smirnov (KS) test. Cell counts for lineage tracing, node / paranode length, and proportion of myelinated axons are presented as mean per mouse ± SD and analysed using a 2-way ANOVA with Sidak post-test multiple comparisons. Behavioural data collected over multiple time points are presented as mean ± SEM and analysed using a mixed two-way ANOVA (sphericity was not assumed), followed by a Sidak-corrected post-test. Dendritic spine densities were analysed by 3-way mixed ANOVA that included genotype as a between-subject factor, and dendrite (apical, basal) and spine morphological type (thin, mushroom, stubby) as repeated measures factors. Significant interactions were followed up with Bonferroni-corrected simple effects tests comparing genotype. The average proportion of spines of any morphology are presented as median ± 95% confidence intervals and were analysed using a Mann-Whitney U for two-group comparisons or Kruskal-Wallis test for independent samples (KW) as noted in each figure legend. ANOVA main effects and interactions are reported in the corresponding figure legends. Statistical significance was defined as p<0.05.

## Notes

### Competing Interest Statement

The authors have declared no competing interest.

## References

1. Tomassy, G.S., Berger, D.R., Chen, H.H., Kasthuri, N., Hayworth, K.J., Vercelli, A., Seung, H.S., Lichtman, J.W., and Arlotta, P. (2014). Distinct profiles of myelin distribution along single axons of pyramidal neurons in the neocortex. Science 344, 319–324. 10.1126/science.1249766.

2. Young, K.M., Psachoulia, K., Tripathi, R.B., Dunn, S.J., Cossell, L., Attwell, D., Tohyama, K., and Richardson, W.D. (2013). Oligodendrocyte dynamics in the healthy adult CNS: evidence for myelin remodeling. Neuron 77, 873–885. 10.1016/j.neuron.2013.01.006.

3. Hughes, E.G., Orthmann-Murphy, J.L., Langseth, A.J., and Bergles, D.E. (2018). Myelin remodeling through experience-dependent oligodendrogenesis in the adult somatosensory cortex. Nat Neurosci 21, 696–706. 10.1038/s41593-018-0121-5.

4. Micheva, K.D., Wolman, D., Mensh, B.D., Pax, E., Buchanan, J., Smith, S.J., and Bock, D.D. (2016). A large fraction of neocortical myelin ensheathes axons of local inhibitory neurons. Elife 5. 10.7554/eLife.15784.

5. Stedehouder, J., Brizee, D., Shpak, G., and Kushner, S.A. (2018). Activity-Dependent Myelination of Parvalbumin Interneurons Mediated by Axonal Morphological Plasticity. J Neurosci 38, 3631–3642. 10.1523/JNEUROSCI.0074-18.2018.

6. Stedehouder, J., Couey, J.J., Brizee, D., Hosseini, B., Slotman, J.A., Dirven, C.M.F., Shpak, G., Houtsmuller, A.B., and Kushner, S.A. (2017). Fast-spiking Parvalbumin Interneurons are Frequently Myelinated in the Cerebral Cortex of Mice and Humans. Cereb Cortex 27, 5001–5013. 10.1093/cercor/bhx203.

7. McKenzie, I.A., Ohayon, D., Li, H., de Faria, J.P., Emery, B., Tohyama, K., and Richardson, W.D. (2014). Motor skill learning requires active central myelination. Science 346, 318–322. 10.1126/science.1254960.

8. Gibson, E.M., Purger, D., Mount, C.W., Goldstein, A.K., Lin, G.L., Wood, L.S., Inema, I., Miller, S.E., Bieri, G., Zuchero, J.B., et al. (2014). Neuronal activity promotes oligodendrogenesis and adaptive myelination in the mammalian brain. Science 344, 1252304. 10.1126/science.1252304.

9. Bacmeister, C.M., Barr, H.J., McClain, C.R., Thornton, M.A., Nettles, D., Welle, C.G., and Hughes, E.G. (2020). Motor learning promotes remyelination via new and surviving oligodendrocytes. Nat Neurosci 23, 819–831. 10.1038/s41593-020-0637-3.

10. Xiao, L., Ohayon, D., McKenzie, I.A., Sinclair-Wilson, A., Wright, J.L., Fudge, A.D., Emery, B., Li, H., and Richardson, W.D. (2016). Rapid production of new oligodendrocytes is required in the earliest stages of motor-skill learning. Nat Neurosci 19, 1210–1217. 10.1038/nn.4351.

11. Barrett, E.F., and Barrett, J.N. (1982). Intracellular recording from vertebrate myelinated axons: mechanism of the depolarizing afterpotential. J Physiol 323, 117–144. 10.1113/jphysiol.1982.sp014064.

12. Freeman, S.A., Desmazieres, A., Fricker, D., Lubetzki, C., and Sol-Foulon, N. (2016). Mechanisms of sodium channel clustering and its influence on axonal impulse conduction. Cell Mol Life Sci 73, 723–735. 10.1007/s00018-015-2081-1.

13. Lubetzki, C., Zalc, B., Williams, A., Stadelmann, C., and Stankoff, B. (2020). Remyelination in multiple sclerosis: from basic science to clinical translation. Lancet Neurol 19, 678–688. 10.1016/S1474-4422(20)30140-X.

14. Saab, A.S., Tzvetavona, I.D., Trevisiol, A., Baltan, S., Dibaj, P., Kusch, K., Mobius, W., Goetze, B., Jahn, H.M., Huang, W., et al. (2016). Oligodendroglial NMDA Receptors Regulate Glucose Import and Axonal Energy Metabolism. Neuron 91, 119–132. 10.1016/j.neuron.2016.05.016.

15. Meyer, N., Richter, N., Fan, Z., Siemonsmeier, G., Pivneva, T., Jordan, P., Steinhauser, C., Semtner, M., Nolte, C., and Kettenmann, H. (2018). Oligodendrocytes in the Mouse Corpus Callosum Maintain Axonal Function by Delivery of Glucose. Cell Rep 22, 2383–2394. 10.1016/j.celrep.2018.02.022.

16. Brown, A.M., Tekkok, S.B., and Ransom, B.R. (2003). Glycogen regulation and functional role in mouse white matter. J Physiol 549, 501–512. 10.1113/jphysiol.2003.042416.

17. Fruhbeis, C., Kuo-Elsner, W.P., Muller, C., Barth, K., Peris, L., Tenzer, S., Mobius, W., Werner, H.B., Nave, K.A., Frohlich, D., and Kramer-Albers, E.M. (2020). Oligodendrocytes support axonal transport and maintenance via exosome secretion. PLoS Biol 18, e3000621. 10.1371/journal.pbio.3000621.

18. Reh, R.K., Dias, B.G., Nelson, C.A., 3rd, Kaufer, D., Werker, J.F., Kolb, B., Levine, J.D., and Hensch, T.K. (2020). Critical period regulation across multiple timescales. Proc Natl Acad Sci U S A 117, 23242–23251. 10.1073/pnas.1820836117.

19. Fletcher, J.L., Makowiecki, K., Cullen, C.L., and Young, K.M. (2021). Oligodendrogenesis and myelination regulate cortical development, plasticity and circuit function. Semin Cell Dev Biol. 10.1016/j.semcdb.2021.03.017.

20. Boghdadi, A.G., Teo, L., and Bourne, J.A. (2018). The Involvement of the Myelin-Associated Inhibitors and Their Receptors in CNS Plasticity and Injury. Mol Neurobiol 55, 1831–1846. 10.1007/s12035-017-0433-6.

21. Roy, K., Murtie, J.C., El-Khodor, B.F., Edgar, N., Sardi, S.P., Hooks, B.M., Benoit-Marand, M., Chen, C., Moore, H., O’Donnell, P., et al. (2007). Loss of erbB signaling in oligodendrocytes alters myelin and dopaminergic function, a potential mechanism for neuropsychiatric disorders. Proc Natl Acad Sci U S A 104, 8131–8136. 10.1073/pnas.0702157104.

22. Chen, X., Zhang, W., Li, T., Guo, Y., Tian, Y., Wang, F., Liu, S., Shen, H.Y., Feng, Y., and Xiao, L. (2015). Impairment of Oligodendroglia Maturation Leads to Aberrantly Increased Cortical Glutamate and Anxiety-Like Behaviors in Juvenile Mice. Front Cell Neurosci 9, 467. 10.3389/fncel.2015.00467.

23. Maheras, K.J., Peppi, M., Ghoddoussi, F., Galloway, M.P., Perrine, S.A., and Gow, A. (2018). Absence of Claudin 11 in CNS Myelin Perturbs Behavior and Neurotransmitter Levels in Mice. Sci Rep 8, 3798. 10.1038/s41598-018-22047-9.

24. Zemmar, A., Chen, C.-C., Weinmann, O., Kast, B., Vajda, F., Bozeman, J., Isaad, N., Zuo, Y., and Schwab, M.E. (2018). Oligodendrocyte- and Neuron-Specific Nogo-A Restrict Dendritic Branching and Spine Density in the Adult Mouse Motor Cortex. Cerebral cortex (New York, N.Y. : 1991) *28*, 2109–2117. 10.1093/cercor/bhx116.

25. Xin, W., Kaneko, M., Roth, R.H., Zhang, A., Nocera, S., Ding, J.B., Stryker, M.P., and Chan, J.R. (2024). Oligodendrocytes and myelin limit neuronal plasticity in visual cortex. Nature 633, 856–863. 10.1038/s41586-024-07853-8.

26. Hamada, M.S., and Kole, M.H. (2015). Myelin loss and axonal ion channel adaptations associated with gray matter neuronal hyperexcitability. J Neurosci 35, 7272–7286. 10.1523/JNEUROSCI.4747-14.2015.

27. Tripathi, R.B., Jackiewicz, M., McKenzie, I.A., Kougioumtzidou, E., Grist, M., and Richardson, W.D. (2017). Remarkable Stability of Myelinating Oligodendrocytes in Mice. Cell Rep 21, 316–323. 10.1016/j.celrep.2017.09.050.

28. Yeung, M.S., Zdunek, S., Bergmann, O., Bernard, S., Salehpour, M., Alkass, K., Perl, S., Tisdale, J., Possnert, G., Brundin, L., et al. (2014). Dynamics of oligodendrocyte generation and myelination in the human brain. Cell 159, 766–774. 10.1016/j.cell.2014.10.011.

29. Dimou, L., Simon, C., Kirchhoff, F., Takebayashi, H., and Gotz, M. (2008). Progeny of Olig2-expressing progenitors in the gray and white matter of the adult mouse cerebral cortex. J Neurosci 28, 10434–10442. 10.1523/JNEUROSCI.2831-08.2008.

30. Rivers, L.E., Young, K.M., Rizzi, M., Jamen, F., Psachoulia, K., Wade, A., Kessaris, N., and Richardson, W.D. (2008). PDGFRA/NG2 glia generate myelinating oligodendrocytes and piriform projection neurons in adult mice. Nat Neurosci 11, 1392–1401. 10.1038/nn.2220.

31. Hill, R.A., Li, A.M., and Grutzendler, J. (2018). Lifelong cortical myelin plasticity and age-related degeneration in the live mammalian brain. Nat Neurosci 21, 683–695. 10.1038/s41593-018-0120-6.

32. Kang, S.H., Fukaya, M., Yang, J.K., Rothstein, J.D., and Bergles, D.E. (2010). NG2+ CNS glial progenitors remain committed to the oligodendrocyte lineage in postnatal life and following neurodegeneration. Neuron 68, 668–681. 10.1016/j.neuron.2010.09.009.

33. Pan, S., Mayoral, S.R., Choi, H.S., Chan, J.R., and Kheirbek, M.A. (2020). Preservation of a remote fear memory requires new myelin formation. Nat Neurosci 23, 487–499. 10.1038/s41593-019-0582-1.

34. Steadman, P.E., Xia, F., Ahmed, M., Mocle, A.J., Penning, A.R.A., Geraghty, A.C., Steenland, H.W., Monje, M., Josselyn, S.A., and Frankland, P.W. (2020). Disruption of Oligodendrogenesis Impairs Memory Consolidation in Adult Mice. Neuron 105, 150–164 e156. 10.1016/j.neuron.2019.10.013.

35. Nabavi, S., Fox, R., Proulx, C.D., Lin, J.Y., Tsien, R.Y., and Malinow, R. (2014). Engineering a memory with LTD and LTP. Nature 511, 348–352. 10.1038/nature13294.

36. Hayashi-Takagi, A., Yagishita, S., Nakamura, M., Shirai, F., Wu, Y.I., Loshbaugh, A.L., Kuhlman, B., Hahn, K.M., and Kasai, H. (2015). Labelling and optical erasure of synaptic memory traces in the motor cortex. Nature 525, 333–338. 10.1038/nature15257.

37. Xu, T., Yu, X., Perlik, A.J., Tobin, W.F., Zweig, J.A., Tennant, K., Jones, T., and Zuo, Y. (2009). Rapid formation and selective stabilization of synapses for enduring motor memories. Nature 462, 915–919. 10.1038/nature08389.

38. Geng, H.Y., Arbuthnott, G., Yung, W.H., and Ke, Y. (2022). Long-range monosynaptic inputs targeting apical and basal dendrites of primary motor cortex deep output neurons. Cereb Cortex 32, 3975–3989. 10.1093/cercor/bhab460.

39. Ramot, A., Taschbach, F.H., Yang, Y.C., Hu, Y., Chen, Q., Morales, B.C., Wang, X.C., Wu, A., Tye, K.M., Benna, M.K., and Komiyama, T. (2025). Motor learning refines thalamic influence on motor cortex. Nature 643, 725–734. 10.1038/s41586-025-08962-8.

40. Neske, G.T., and Cardin, J.A. (2025). Higher-order thalamic input to cortex selectively conveys state information. Cell Rep 44, 115292. 10.1016/j.celrep.2025.115292.

41. Hwang, F.J., Roth, R.H., Wu, Y.W., Sun, Y., Kwon, D.K., Liu, Y., and Ding, J.B. (2022). Motor learning selectively strengthens cortical and striatal synapses of motor engram neurons. Neuron 110, 2790–2801 e2795. 10.1016/j.neuron.2022.06.006.

42. Albarran, E., Raissi, A., Jaidar, O., Shatz, C.J., and Ding, J.B. (2021). Enhancing motor learning by increasing the stability of newly formed dendritic spines in the motor cortex. Neuron 109, 3298–3311 e3294. 10.1016/j.neuron.2021.07.030.

43. Li, Q., Ko, H., Qian, Z.M., Yan, L.Y.C., Chan, D.C.W., Arbuthnott, G., Ke, Y., and Yung, W.H. (2017). Refinement of learned skilled movement representation in motor cortex deep output layer. Nat Commun 8, 15834. 10.1038/ncomms15834.

44. Kida, H., Kawakami, R., Sakai, K., Otaku, H., Imamura, K., Han, T.Z., Sakimoto, Y., and Mitsushima, D. (2023). Motor training promotes both synaptic and intrinsic plasticity of layer V pyramidal neurons in the primary motor cortex. J Physiol 601, 335–353. 10.1113/JP283755.

45. Kida, H., Toyoshima, S., Kawakami, R., Sakimoto, Y., and Mitsushima, D. (2024). Properties of layer V pyramidal neurons in the primary motor cortex that represent acquired motor skills. Neuroscience 559, 54–63. 10.1016/j.neuroscience.2024.08.033.

46. Emery, B., Agalliu, D., Cahoy, J.D., Watkins, T.A., Dugas, J.C., Mulinyawe, S.B., Ibrahim, A., Ligon, K.L., Rowitch, D.H., and Barres, B.A. (2009). Myelin gene regulatory factor is a critical transcriptional regulator required for CNS myelination. Cell 138, 172–185. 10.1016/j.cell.2009.04.031.

47. Koenning, M., Jackson, S., Hay, C.M., Faux, C., Kilpatrick, T.J., Willingham, M., and Emery, B. (2012). Myelin gene regulatory factor is required for maintenance of myelin and mature oligodendrocyte identity in the adult CNS. J Neurosci 32, 12528–12542. 10.1523/JNEUROSCI.1069-12.2012.

48. Watanabe, M., Toyama, Y., and Nishiyama, A. (2002). Differentiation of proliferated NG2-positive glial progenitor cells in a remyelinating lesion. J Neurosci Res 69, 826–836. 10.1002/jnr.10338.

49. Cullen, C.L., Pepper, R.E., Clutterbuck, M.T., Pitman, K.A., Oorschot, V., Auderset, L., Tang, A.D., Ramm, G., Emery, B., Rodger, J., et al. (2021). Periaxonal and nodal plasticities modulate action potential conduction in the adult mouse brain. Cell Reports 34, 108641. 10.1016/j.celrep.2020.108641.

50. Clark, J.A., Blizzard, C.A., Breslin, M.C., Yeaman, E.J., Lee, K.M., Chuckowree, J.A., and Dickson, T.C. (2018). Epothilone D accelerates disease progression in the SOD1(G93A) mouse model of amyotrophic lateral sclerosis. Neuropathol Appl Neurobiol 44, 590–605. 10.1111/nan.12473.

51. Duncan, G.J., Ingram, S.D., Emberley, K., Hill, J., Cordano, C., Abdelhak, A., McCane, M., Jenks, J.E., Jabassini, N., Ananth, K., et al. (2024). Remyelination protects neurons from DLK-mediated neurodegeneration. Nat Commun 15, 9148. 10.1038/s41467-024-53429-5.

52. Spruston, N. (2008). Pyramidal neurons: dendritic structure and synaptic integration. Nat Rev Neurosci 9, 206–221. 10.1038/nrn2286.

53. Oswald, M.J., Tantirigama, M.L., Sonntag, I., Hughes, S.M., and Empson, R.M. (2013). Diversity of layer 5 projection neurons in the mouse motor cortex. Front Cell Neurosci 7, 174. 10.3389/fncel.2013.00174.

54. Wright, W.J., Hedrick, N.G., and Komiyama, T. (2025). Distinct synaptic plasticity rules operate across dendritic compartments in vivo during learning. Science 388, 322–328. 10.1126/science.ads4706.

55. Matsuzaki, M., Honkura, N., Ellis-Davies, G.C., and Kasai, H. (2004). Structural basis of long-term potentiation in single dendritic spines. Nature 429, 761–766. 10.1038/nature02617.

56. Holtmaat, A.J., Trachtenberg, J.T., Wilbrecht, L., Shepherd, G.M., Zhang, X., Knott, G.W., and Svoboda, K. (2005). Transient and persistent dendritic spines in the neocortex in vivo. Neuron 45, 279–291. 10.1016/j.neuron.2005.01.003.

57. Bailey, C.H., Kandel, E.R., and Harris, K.M. (2015). Structural Components of Synaptic Plasticity and Memory Consolidation. Cold Spring Harb Perspect Biol 7, a021758. 10.1101/cshperspect.a021758.

58. Young, K.M., Mitsumori, T., Pringle, N., Grist, M., Kessaris, N., and Richardson, W.D. (2010). An Fgfr3-iCreER(T2) transgenic mouse line for studies of neural stem cells and astrocytes. Glia 58, 943–953. 10.1002/glia.20976.

59. Xing, Y.L., Roth, P.T., Stratton, J.A., Chuang, B.H., Danne, J., Ellis, S.L., Ng, S.W., Kilpatrick, T.J., and Merson, T.D. (2014). Adult neural precursor cells from the subventricular zone contribute significantly to oligodendrocyte regeneration and remyelination. J Neurosci 34, 14128–14146. 10.1523/JNEUROSCI.3491-13.2014.

60. Zhi, J.J., Wu, S.L., Wu, H.Q., Ran, Q., Gao, X., Chen, J.F., Gu, X.M., Li, T., Wang, F., Xiao, L., et al. (2023). Insufficient Oligodendrocyte Turnover in Optic Nerve Contributes to Age-Related Axon Loss and Visual Deficits. J Neurosci 43, 1859–1870. 10.1523/JNEUROSCI.2130-22.2023.

61. Schneider, S., Gruart, A., Grade, S., Zhang, Y., Kroger, S., Kirchhoff, F., Eichele, G., Delgado Garcia, J.M., and Dimou, L. (2016). Decrease in newly generated oligodendrocytes leads to motor dysfunctions and changed myelin structures that can be rescued by transplanted cells. Glia 64, 2201–2218. 10.1002/glia.23055.

62. Cullen, C.L., O’Rourke, M., Beasley, S.J., Auderset, L., Zhen, Y., Pepper, R.E., Gasperini, R., and Young, K.M. (2020). Kif3a deletion prevents primary cilia assembly on oligodendrocyte progenitor cells, reduces oligodendrogenesis and impairs fine motor function. Glia. 10.1002/glia.23957.

63. Quinn, D.P., Kolar, A., Harris, S.A., Wigerius, M., Fawcett, J.P., and Krueger, S.R. (2019). The Stability of Glutamatergic Synapses Is Independent of Activity Level, but Predicted by Synapse Size. Front Cell Neurosci 13, 291. 10.3389/fncel.2019.00291.

64. Nguyen, P.T., Makowiecki, K., Lewis, T.S., Fortune, A.J., Clutterbuck, M., Reale, L.A., Taylor, B.V., Rodger, J., Cullen, C.L., and Young, K.M. (2024). Low intensity repetitive transcranial magnetic stimulation enhances remyelination by newborn and surviving oligodendrocytes in the cuprizone model of toxic demyelination. Cell Mol Life Sci 81, 346. 10.1007/s00018-024-05391-0.

65. Guadalupi, L., Mandolesi, G., Vanni, V., Balletta, S., Caioli, S., Pavlovic, A., De Vito, F., Fresegna, D., Sanna, K., Vitiello, L., et al. (2024). Pharmacological blockade of 2-AG degradation ameliorates clinical, neuroinflammatory and synaptic alterations in experimental autoimmune encephalomyelitis. Neuropharmacology 252, 109940. 10.1016/j.neuropharm.2024.109940.

66. Hobbiss, A.F., Ramiro-Cortes, Y., and Israely, I. (2018). Homeostatic Plasticity Scales Dendritic Spine Volumes and Changes the Threshold and Specificity of Hebbian Plasticity. iScience 8, 161–174. 10.1016/j.isci.2018.09.015.

67. Hu, H., Real, E., Takamiya, K., Kang, M.G., Ledoux, J., Huganir, R.L., and Malinow, R. (2007). Emotion enhances learning via norepinephrine regulation of AMPA-receptor trafficking. Cell 131, 160–173. 10.1016/j.cell.2007.09.017.

68. Guo, L., Xiong, H., Kim, J.I., Wu, Y.W., Lalchandani, R.R., Cui, Y., Shu, Y., Xu, T., and Ding, J.B. (2015). Dynamic rewiring of neural circuits in the motor cortex in mouse models of Parkinson’s disease. Nat Neurosci 18, 1299–1309. 10.1038/nn.4082.

69. Macrez, R., Stys, P.K., Vivien, D., Lipton, S.A., and Docagne, F. (2016). Mechanisms of glutamate toxicity in multiple sclerosis: biomarker and therapeutic opportunities. Lancet Neurol 15, 1089–1102. 10.1016/S1474-4422(16)30165-X.

70. Huiskamp, M., Kiljan, S., Kulik, S., Witte, M.E., Jonkman, L.E., Gjm Bol, J., Schenk, G.J., Hulst, H.E., Tewarie, P., Schoonheim, M.M., and Geurts, J.J. (2022). Inhibitory synaptic loss drives network changes in multiple sclerosis: An ex vivo to in silico translational study. Mult Scler 28, 2010–2019. 10.1177/13524585221125381.

71. Jurgens, T., Jafari, M., Kreutzfeldt, M., Bahn, E., Bruck, W., Kerschensteiner, M., and Merkler, D. (2016). Reconstruction of single cortical projection neurons reveals primary spine loss in multiple sclerosis. Brain 139, 39–46. 10.1093/brain/awv353.

72. Dutta, R., Chang, A., Doud, M.K., Kidd, G.J., Ribaudo, M.V., Young, E.A., Fox, R.J., Staugaitis, S.M., and Trapp, B.D. (2011). Demyelination causes synaptic alterations in hippocampi from multiple sclerosis patients. Ann Neurol 69, 445–454. 10.1002/ana.22337.

73. Mock, E.E.A., Honkonen, E., and Airas, L. (2021). Synaptic Loss in Multiple Sclerosis: A Systematic Review of Human Post-mortem Studies. Front Neurol 12, 782599. 10.3389/fneur.2021.782599.

74. Cordano, C., Werneburg, S., Abdelhak, A., Bennett, D.J., Beaudry-Richard, A., Duncan, G.J., Oertel, F.C., Boscardin, W.J., Yiu, H.H., Jabassini, N., et al. (2024). Synaptic injury in the inner plexiform layer of the retina is associated with progression in multiple sclerosis. Cell Rep Med 5, 101490. 10.1016/j.xcrm.2024.101490.

75. Werneburg, S., Jung, J., Kunjamma, R.B., Ha, S.K., Luciano, N.J., Willis, C.M., Gao, G., Biscola, N.P., Havton, L.A., Crocker, S.J., et al. (2020). Targeted Complement Inhibition at Synapses Prevents Microglial Synaptic Engulfment and Synapse Loss in Demyelinating Disease. Immunity 52, 167–182 e167. 10.1016/j.immuni.2019.12.004.

76. Gillani, R.L., Kironde, E.N., Whiteman, S., Zwang, T.J., and Bacskai, B.J. (2024). Instability of excitatory synapses in experimental autoimmune encephalomyelitis and the outcome for excitatory circuit inputs to individual cortical neurons. Brain Behav Immun 119, 251–260. 10.1016/j.bbi.2024.03.039.

77. Potter, L.E., Paylor, J.W., Suh, J.S., Tenorio, G., Caliaperumal, J., Colbourne, F., Baker, G., Winship, I., and Kerr, B.J. (2016). Altered excitatory-inhibitory balance within somatosensory cortex is associated with enhanced plasticity and pain sensitivity in a mouse model of multiple sclerosis. J Neuroinflammation 13, 142. 10.1186/s12974-016-0609-4.

78. Lohrberg, M., Mortensen, L.S., Thomas, C., Fries, F., van der Meer, F., Gotz, A., Landt, C., Rhee, H.J., Rhee, J., Gomez-Varela, D., et al. (2025). Astroglial modulation of synaptic function in the non-demyelinated cerebellar cortex is dependent on MyD88 signaling in a model of toxic demyelination. J Neuroinflammation 22, 47. 10.1186/s12974-025-03368-9.

79. Baltan, S., Jawaid, S.S., Chomyk, A.M., Kidd, G.J., Chen, J., Battapady, H.D., Chan, R., Dutta, R., and Trapp, B.D. (2021). Neuronal hibernation following hippocampal demyelination. Acta Neuropathol Commun 9, 34. 10.1186/s40478-021-01130-9.

80. Trachtenberg, J.T., Chen, B.E., Knott, G.W., Feng, G., Sanes, J.R., Welker, E., and Svoboda, K. (2002). Long-term in vivo imaging of experience-dependent synaptic plasticity in adult cortex. Nature 420, 788–794. 10.1038/nature01273.

81. Grutzendler, J., Kasthuri, N., and Gan, W.B. (2002). Long-term dendritic spine stability in the adult cortex. Nature 420, 812–816. 10.1038/nature01276.

82. Zuo, Y., Lin, A., Chang, P., and Gan, W.B. (2005). Development of long-term dendritic spine stability in diverse regions of cerebral cortex. Neuron 46, 181–189. 10.1016/j.neuron.2005.04.001.

83. Munoz-Castaneda, R., Zingg, B., Matho, K.S., Chen, X., Wang, Q., Foster, N.N., Li, A., Narasimhan, A., Hirokawa, K.E., Huo, B., et al. (2021). Cellular anatomy of the mouse primary motor cortex. Nature 598, 159–166. 10.1038/s41586-021-03970-w.

84. Klionsky, D.J., Abdelmohsen, K., Abe, A., Abedin, M.J., Abeliovich, H., Acevedo Arozena, A., Adachi, H., Adams, C.M., Adams, P.D., Adeli, K., et al. (2016). Guidelines for the use and interpretation of assays for monitoring autophagy (3rd edition). Autophagy 12, 1–222. 10.1080/15548627.2015.1100356.

85. Rioult-Pedotti, M.S., Friedman, D., and Donoghue, J.P. (2000). Learning-induced LTP in neocortex. Science 290, 533–536. 10.1126/science.290.5491.533.

86. Roth, R.H., Cudmore, R.H., Tan, H.L., Hong, I., Zhang, Y., and Huganir, R.L. (2020). Cortical Synaptic AMPA Receptor Plasticity during Motor Learning. Neuron 105, 895–908 e895. 10.1016/j.neuron.2019.12.005.

87. Doerflinger, N.H., Macklin, W.B., and Popko, B. (2003). Inducible site-specific recombination in myelinating cells. Genesis 35, 63–72. 10.1002/gene.10154.

88. Srinivas, S., Watanabe, T., Lin, C.S., William, C.M., Tanabe, Y., Jessell, T.M., and Costantini, F. (2001). Cre reporter strains produced by targeted insertion of EYFP and ECFP into the ROSA26 locus. BMC Dev Biol 1, 4. 10.1186/1471-213x-1-4.

89. Feng, G., Mellor, R.H., Bernstein, M., Keller-Peck, C., Nguyen, Q.T., Wallace, M., Nerbonne, J.M., Lichtman, J.W., and Sanes, J.R. (2000). Imaging neuronal subsets in transgenic mice expressing multiple spectral variants of GFP. Neuron 28, 41–51. 10.1016/s0896-6273(00)00084-2.

90. Auderset, L., Pitman, K.A., Cullen, C.L., Pepper, R.E., Taylor, B.V., Foa, L., and Young, K.M. (2020). Low-Density Lipoprotein Receptor-Related Protein 1 (LRP1) Is a Negative Regulator of Oligodendrocyte Progenitor Cell Differentiation in the Adult Mouse Brain. Front Cell Dev Biol 8, 564351. 10.3389/fcell.2020.564351.

91. Cashion, J.M., Brown, L.S., Morris, G.P., Fortune, A.J., Courtney, J.M., Makowiecki, K., Premilovac, D., Cullen, C.L., Young, K.M., and Sutherland, B.A. (2025). Pericyte ablation causes hypoactivity and reactive gliosis in adult mice. Brain Behav Immun 123, 681–696. 10.1016/j.bbi.2024.10.014.

92. Handley, E.E., Pitman, K.A., Dawkins, E., Young, K.M., Clark, R.M., Jiang, T.C., Turner, B.J., Dickson, T.C., and Blizzard, C.A. (2017). Synapse Dysfunction of Layer V Pyramidal Neurons Precedes Neurodegeneration in a Mouse Model of TDP-43 Proteinopathies. Cereb Cortex 27, 3630–3647. 10.1093/cercor/bhw185.

