## Supplementary Figures for "Myelin maintenance and addition regulate synaptic plasticity in the adult mouse cortex"

Supplementary Data Pepper et al.

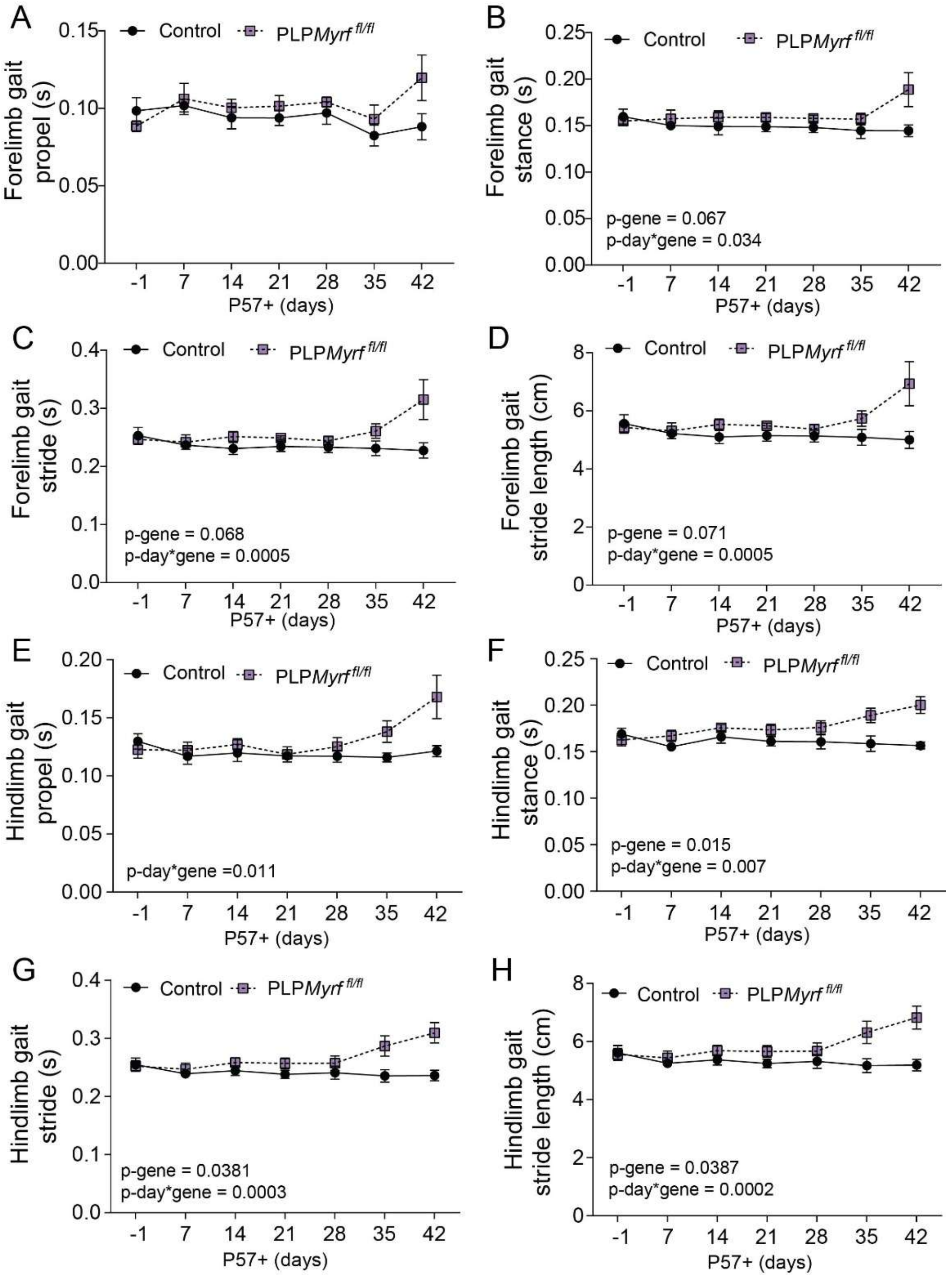


**Figure S1:** The conditional deletion of Myrf from OLs at ~2 months of age alters running gait.

A-C) Average forelimb propel time (A), stance time (B) and stride time (C) during one full step by control (black circles) and PLP*Myrf* $^{fl/fl}$ (purple squares) mice during treadmill (DigiGait™) running from P57-1 to P57+42. [Restricted Maximum Likelihood (REML) mixed effects model: *Propel time* (Interaction F(6, 53)=1.602 p=0.16; day F(6, 53)=1.58 p=0.17; genotype F(1,13)=2.48 p=0.13); *Stance time* (Interaction F(6, 53)=2.48 p=0.034; day F(2.738, 24.18)=1.328 p=0.28; genotype

F(1,13)=3.995 p=0.067); *Stride time* (Interaction F(6, 53)=4.87 p=0.0005; day F(2.274, 20.09)=2.121 p=0.14; genotype F(1,13)=3.933 p=0.068)]

D) Average forelimb stride length during one full step by control (black circles) and PLP*Myrf* $^{fl/fl}$ (purple squares) mice during treadmill (DigiGait$^{TM}$) running from P57-1 to P57+42. [Restricted Maximum Likelihood (REML) mixed effects model: Interaction F(6, 53)=4.91 p=0.0005; day F(2.316, 20.46)=2.076 p=0.14; genotype F(1,13)=3.855 p=0.071]

E-G) Average hindlimb propel time (E), stance time (F) and stride time (G) during one full step by control (black circles) and PLP*Myrf* $^{fl/fl}$ (purple squares) mice during treadmill (DigiGait$^{TM}$) running from P57-1 to P57+42. [Restricted Maximum Likelihood (REML) mixed effects model: *Propel time* (Interaction F(6, 53)=3.099 p=0.011; day F(2.422, 21.40)=3.07 p=0.058; genotype F(1,13)=2.84 p=0.11); *Stance time* (Interaction F(6, 53)=3.321 p=0.007; day F(3.328, 29.40)=1.449 p=0.24; genotype F(1,13)=7.68 p=0.015); *Stride time* (Interaction F(6, 53)=5.103 p=0.0003; day F(2.775, 24.51)=1.931 p=0.15; genotype F(1,13)=5.328 p=0.038)]

H) Average hindlimb stride length during one full step by control (black circles) and PLP*Myrf* $^{fl/fl}$ (purple squares) mice during treadmill (DigiGait$^{TM}$) running from P57-1 to P57+42. [Restricted Maximum Likelihood (REML) mixed effects model: Interaction F(6, 53)=5.311 p=0.0002; day F(2.697, 23.82)=1.90 p=0.16; genotype F(1,13)=5.285 p=0.038]

Graphs show mean ± SEM. Control n= 6-9; PLPMyrf $^{fl/fl}$ n= 4-7. See also Figure 2

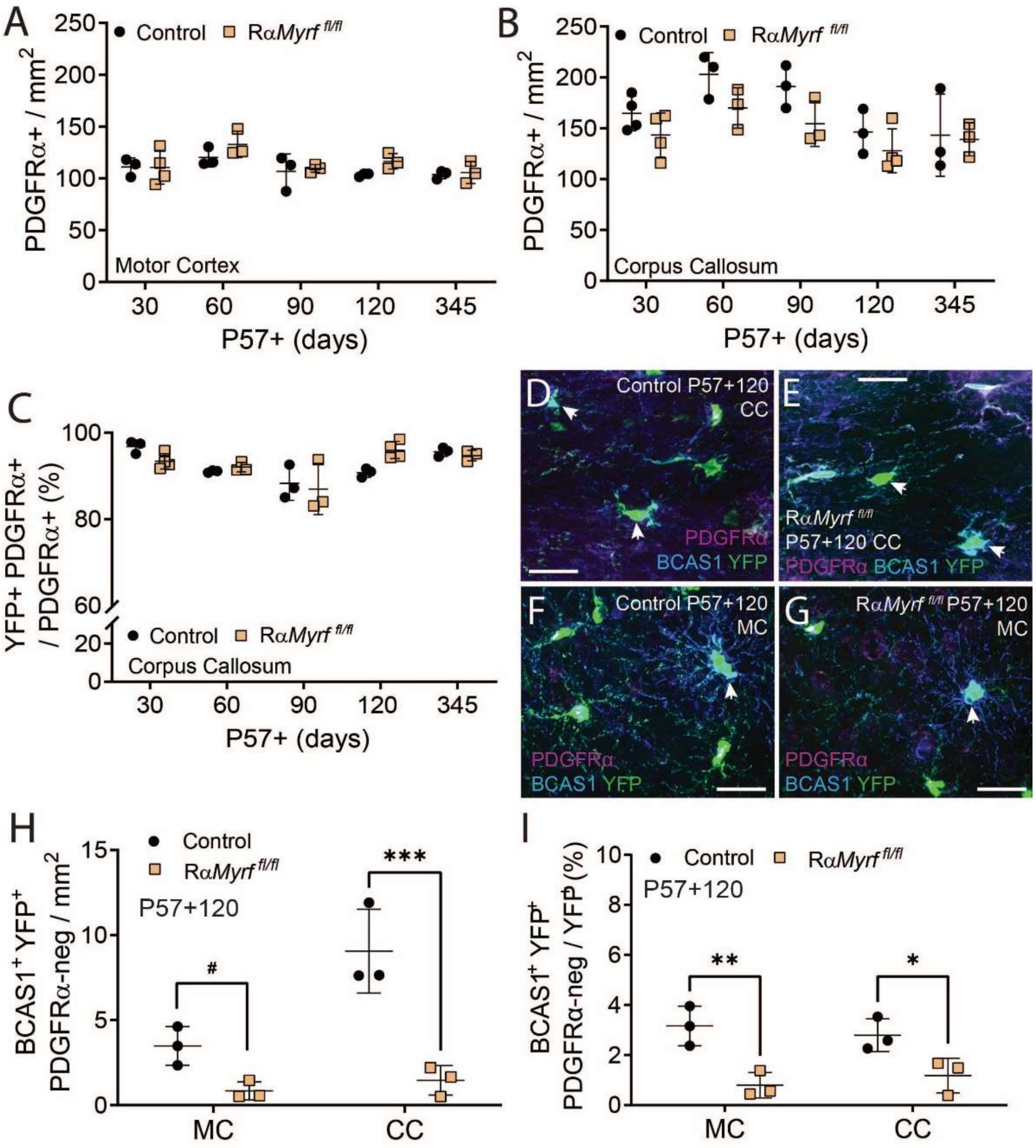


**Figure S2:** Conditionally deleting Myrf from OPCs at ~2 months of age, reduces premyelinating OL formation

A-B) Density of PDGFRα+ OPCs in the motor cortex (A) and corpus callosum (B) of control (black circles) and Rα*Myrf*$^{fl/fl}$ (orange squares) mice at P57+30 to P57+345 days. Two-way ANOVA: *motor cortex* Interaction F(4, 21)=0.544 p=0.704; day F(4, 21)=3.71 p=0.019; genotype F(1,21)=2.45 p=0.13; *corpus callosum* Interaction F(4, 23)=0.478 p=0.75; day F(4, 23)=5.23 p=0.003; genotype F(1,23)=7.931 p=0.009]

C) Proportion of recombined OPCs (YFP$^{+}$ PDGFRα$^{+}$) in the corpus callosum of of control (black circles) and Rα*Myrf* $^{fl/fl}$ (orange squares) mice from P57+30 to P57+345. Two-way ANOVA: Interaction F(4, 22)=2.893 p=0.045; day F(4, 22)=9.689 p=0.0001; genotype F(1,22)=0.018 p=0.894

D-G) Confocal images from the corpus callosum (CC) and motor cortex (MC) of P57+120 control and Rα*Myrf* $^{fl/fl}$ mice, stained to detect OPCs (PDGFRα, magenta), premyelinating OLs (BCAS1, blue) and YFP (green).

H) Density of BCAS1$^{+}$ YFP$^{+}$ premyelinating OLs (PDGFRα-neg) in the motor cortex (MC) and corpus callosum (CC) of control (black circles) and Rα*Myrf* $^{fl/fl}$ (orange squares) mice at P57+120. Two-way ANOVA: Interaction F(1, 8)=8.75 p=0.018; region F(1, 8)=13.67 p=0.0061; genotype F(1,8)=37.38 p=0.0003.

I) Proportion of total YFP$^{+}$ cells that were BCAS1$^{+}$ premyelinating OLs (PDGFRα-neg) in the motor cortex (MC) and corpus callosum (CC) of control (black circles) and Rα*Myrf* $^{fl/fl}$ (orange squares) mice at P57+120. Two-way ANOVA: Interaction F(1, 8)=0.96 p=0.35; region F(1, 8)=0.0003 p=0.98; genotype F(1,8)=26.50 p=0.0009.

Graphs show mean ± SD, *p<0.05, **p<0.01, ***p<0.001, by Sidak post-test. See also Figure 5.

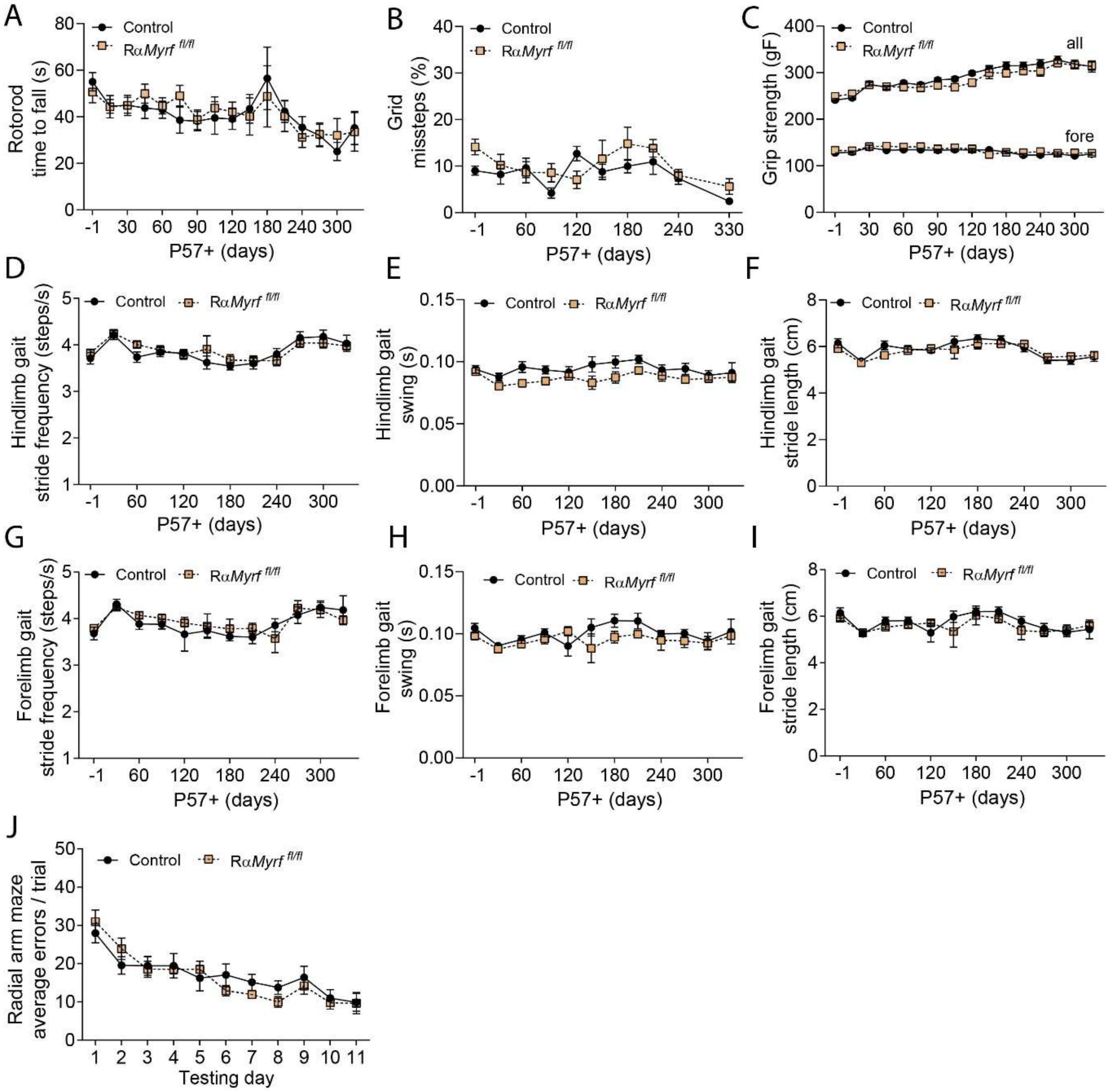


**Figure S3:** Motor function remains largely normal in RαMyrf $^{fl/fl}$ mice

A) Time taken for control (black circles, n= 13-58) and Rα*Myrf*$^{fl/fl}$ (orange squares, n= 9-51) mice to fall from the rotarod at P57-1 to P57+330. [Restricted Maximum Likelihood (REML) mixed effects model: Interaction $F(15, 722) = 0.488$ $p=0.94$; day $F(15, 722) = 3.94$ $p<0.0001$; genotype $F(1, 107) = 0.39$ $p=0.53$].

B) Proportion of missteps made by control (n=9-15) and Rα*Myrf*$^{fl/fl}$ (n=7-9) mice performing the grid walk test at P57-1 to P57+330. [Restricted Maximum Likelihood (REML) mixed effects model: Interaction $F(9, 173) = 1.417$ $p=0.18$; day $F(4.46, 85.75) =3.677$ $p=0.0063$; genotype $F(1, 22) = 2.601$ $p=0.12$].

C) Grip strength for all paws and forepaws only of control (n= 13-58) and Rα*Myrf*$^{fl/fl}$ (n= 9-51) mice from P57-1 to P57+330. [Restricted Maximum Likelihood (REML) mixed effects model: *All paws* (Interaction $F(15, 778) = 2.818$ $p= 0.00033$; day $F(15, 778) = 30.74$ $p<0.0001$; genotype $F(1,122) =$

3.267 p=0.073); *Fore paws* (Interaction F(15, 758)=0.56 p=0.90; day F(15, 758)=3.469 p<0.0001; genotype F(1,112) = 1.057 p=0.306)]

D-E) Average hindlimb stride frequency (D) and limb swing time (E) of control (n=5-18) and Rα*Myrf* $^{fl/fl}$ (n=7-22) mice during treadmill (DigiGait$^{TM}$) running from P57-1 to P57+330. [Restricted Maximum Likelihood (REML) mixed effects model: *Stride frequency* (Interaction F(11, 186) = 0.599 p=0.82; day F(4.432, 74.94) = 8.489 p<0.0001; genotype F(1, 44) = 0.577 p=0.45); *Swing* (Interaction F(11, 187) = 0.956 p=0.48; day F(4.97, 84.49) = 2.945 p=0.017; genotype F(1,45) = 8.112 p=0.006)]

F) Average hindlimb stride length during one full step by control and Rα*Myrf* $^{fl/fl}$ mice during treadmill (DigiGait$^{TM}$) running from P57-1 to P57+330. [Restricted Maximum Likelihood (REML) mixed effects model: Interaction F(11, 186) = 0.565 p=0.85; day F(4.407, 74.52) = 8.54 p<0.0001; genotype F(1,44) = 1.019 p=0.31]

G-H) Average forelimb stride frequency (G) and limb swing time (H) of control and Rα*Myrf*$^{fl/fl}$ mice during treadmill (DigiGait$^{TM}$) running from P57-1 to P57+330. [Restricted Maximum Likelihood (REML) mixed effects model: *Stride frequency* (Interaction F(11, 186) = 0.849 p=0.59; day F(5.79, 98.04) = 7.84 p<0.0001; genotype F(1,44) = 0.964 p=0.33); *Swing* (Interaction F(11, 186) =1.21 p=0.28; day F(5.32, 89.48) = 3.24 p=0.0085; genotype F(1,44)=3.775 p=0.058)]

I) Average forelimb stride length during one full step by control and Rα*Myrf*$^{fl/fl}$ mice during treadmill (DigiGait$^{TM}$) running from P57-1 to P57+330. [Restricted Maximum Likelihood (REML) mixed effects model: Interaction F(11, 186) = 0.77 p=0.66; day F(4.587, 78.40) = 4.328 p=0.0004; genotype F(1,44) = 1.055 p=0.309]

J) Quantification of the average number of errors made across 3 trials each day by control (n=8) and Rα*Myrf*$^{fl/fl}$ (n=7) mice learning the RAM task from P57 + 60. [RM two-way ANOVA: Interaction F(10, 130)=1.467, p=0.158; day F(4.876, 63.39) = 24.28, p<0.0001; genotype F(1, 13) = 0.06, p=0.807]

Graphs show mean ± SEM.
